# KCNQ2 Loss-of-Function variants disrupt neuronal maturation via early hyperexcitability followed by maladaptive network remodeling

**DOI:** 10.1101/2025.07.22.664856

**Authors:** N. Dirkx, M. Kaji, E. De Vriendt, G. Carleo, F. Miceli, B. Asselbergh, P. Verstraelen, N. Zonnekein, L. Carotenuto, L.T. Dang, V. Sommers, E. Vlaemynck, L. Lagae, B. Ceulemans, P. De Jonghe, W. H. De Vos, M. Taglialatela, S. Weckhuysen

## Abstract

Loss-of-function (LOF) variants in the potassium channel subunit *KCNQ2* cause a spectrum of neonatal epilepsies from self-limiting familial neonatal epilepsy (SeLFNE) to severe developmental and epileptic encephalopathy (DEE). To dissect the developmental consequences of *KCNQ2* LOF, we conducted a longitudinal and multimodal comparative analysis in a human neuronal model generated from patients with *KCNQ2*-DEE and *KCNQ2*-SeLFNE. *KCNQ2*-LOF induced a biphasic network dysfunction, with early Kv7-driven hyperexcitability rescued by acute Retigabine (RTG) treatment, followed by maladaptive remodeling in the opposite direction. Transcriptomic analysis mirrored this biphasic dynamic trajectory, revealing an initial upregulation followed by a subsequent downregulation of synaptic genes. Structural analysis showed a steeper decline in presynaptic density alongside a distal shift in the axon initial segment (AIS) throughout maturation, and impaired AIS plasticity at later stages. Overall, KCNQ2-LOF disrupts human neuronal maturation through dynamic, biphasic changes in function, gene expression and structure, offering insights into disease mechanisms and therapeutic options.

## Introduction

The *KCNQ2* gene encodes the Kv7.2 subunit of the voltage-gated potassium channel Kv7, which is widely expressed throughout the brain (1). Kv7 channels are primarily composed of Kv7.2 and Kv7.3 subunits (2). These channels are abundantly expressed at the axon initial segment (AIS), nodes of Ranvier, the soma, and presynaptic terminals (3–5), where they generate the M-current, a slowly activating, non-inactivating potassium current, that stabilizes membrane potential and suppresses repetitive firing (2, 6).

Human pathogenic variants in *KCNQ2* cause a spectrum of epileptic and neurodevelopmental disorders, underscoring the importance of Kv7.2 in normal brain function (7–9). Most are heterozygous loss-of-function (LOF) variants that reduce M-current. Dominant negative (DN) variants cause severe neonatal-onset developmental and epileptic encephalopathy (*KCNQ2*-DEE) (9), while milder LOF variants cause self-limited familial neonatal epilepsy (KCNQ2-SeLFNE), characterized by neonatal seizures that resolve with age and typical development (8). Rare gain-of-function (GOF) variants cause developmental delay with or without later onset seizures (*KCNQ2*-GOF)(7).

While traditional anti-seizure medication (ASM) can alleviate seizures, *KCNQ2*-DEE patients invariably suffer from developmental delay. Retigabine (RTG), a selective Kv7 potassium channel opener used as an ASM for focal-onset seizures, showed promise as a targeted therapy for *KCNQ2*-DEE. A retrospective study involving eight *KCNQ2*-DEE patients treated with RTG reported improved developmental outcomes in all patients, supporting its potential to address Kv7 dysfunction (10). Unfortunately, RTG was withdrawn from the market in June 2017, mainly due to side effects related to its chemical instability (11).

A *KCNQ2*-DEE T274M knock-in mouse model recapitulating key clinical features such as spontaneous seizures and memory impairment, revealed a transient reduction in M-current and increased excitability in pyramidal cells (12, 13). While mouse models provide valuable mechanistic insights, species-specific differences in developmental timing and gene expression limit their translational relevance (14–17). Human induced pluripotent stem cell (hiPSC)-derived neurons offer a complementary platform that enables the study of patient-specific genetic variants in a human model. For instance, iPSC-derived neurons with the *KCNQ2*-DEE R581Q variant exhibited accelerated action potential repolarization, larger post-burst afterhyperpolarization, and dyshomeostatic enhancement of calcium activated potassium channels (18). These findings showcase the utility of studying neuronal activity and development in a human *in vitro* setting to provide direct insight into human disease.

Given the diverse clinical phenotypes associated with *KCNQ2* variants, this study aims to provide a detailed analysis of the developmental trajectories caused by *KCNQ2*-LOF variants in a human neuronal model, through comparative transcriptomic, structural and functional profiling. To this end, we generated iPSC-derived glutamatergic cortical iNeurons of three KCNQ2-DEE patients and one KCNQ2-SeLFNE patient. We characterized these lines across early and late maturation, identifying a biphasic pattern of dysfunction: an early phase of hyperexcitability linked to direct Kv7 channel dysfunction as acute RTG treatment effectively reversed the electrophysiological alterations, followed by a later phase characterized by maladaptive changes, in which RTG exacerbated dysfunction. This biphasic trajectory was also reflected in transcriptomic and structural alterations, particularly affecting synaptic signaling. Together, our study provides new insights into the dynamic consequences of *KCNQ2*-LOF during neuronal maturation, with implications for understanding disease mechanisms and optimizing therapeutic strategies.

## Results

### Generation and characterization of KCNQ2-LOF iNeurons

To study *KCNQ2*-related LOF disorders, we generated iPSC-derived cortical neurons using Neurogenin1/2 (NGN) overexpression (iNeurons) (Fig. 1A-B), from three patients with *KCNQ2*-DEE, each carrying a de novo pathogenic heterozygous *KCNQ2* variant (R560W, G290D, A294V), a KCNQ2-SeLFNE patient belonging to a two-generation pedigree carrying a segregating heterozygous variant (K327Q), a healthy sibling of the *KCNQ2*-R560W patient (WT), and an isogenic control of the *KCNQ2*-DEE G290D variant (ISO_G290D). The KCNQ2-DEE patients presented a typical *KCNQ2*-DEE phenotype consisting of neonatal seizures and developmental delay, the SeLFNE family members exhibited neonatal seizures and typical neurodevelopment (Table 1). All *KCNQ2*-DEE variants were previously reported and functional studies have demonstrated a severe LOF effect (9, 19–21). All *KCNQ2* variants were classified as (likely) pathogenic according to ACMG criteria (22). The *KCNQ2* variants and the correction of the G290D variant were confirmed in the iPSC lines via Sanger sequencing (Fig. S1).

**Fig. 1.**
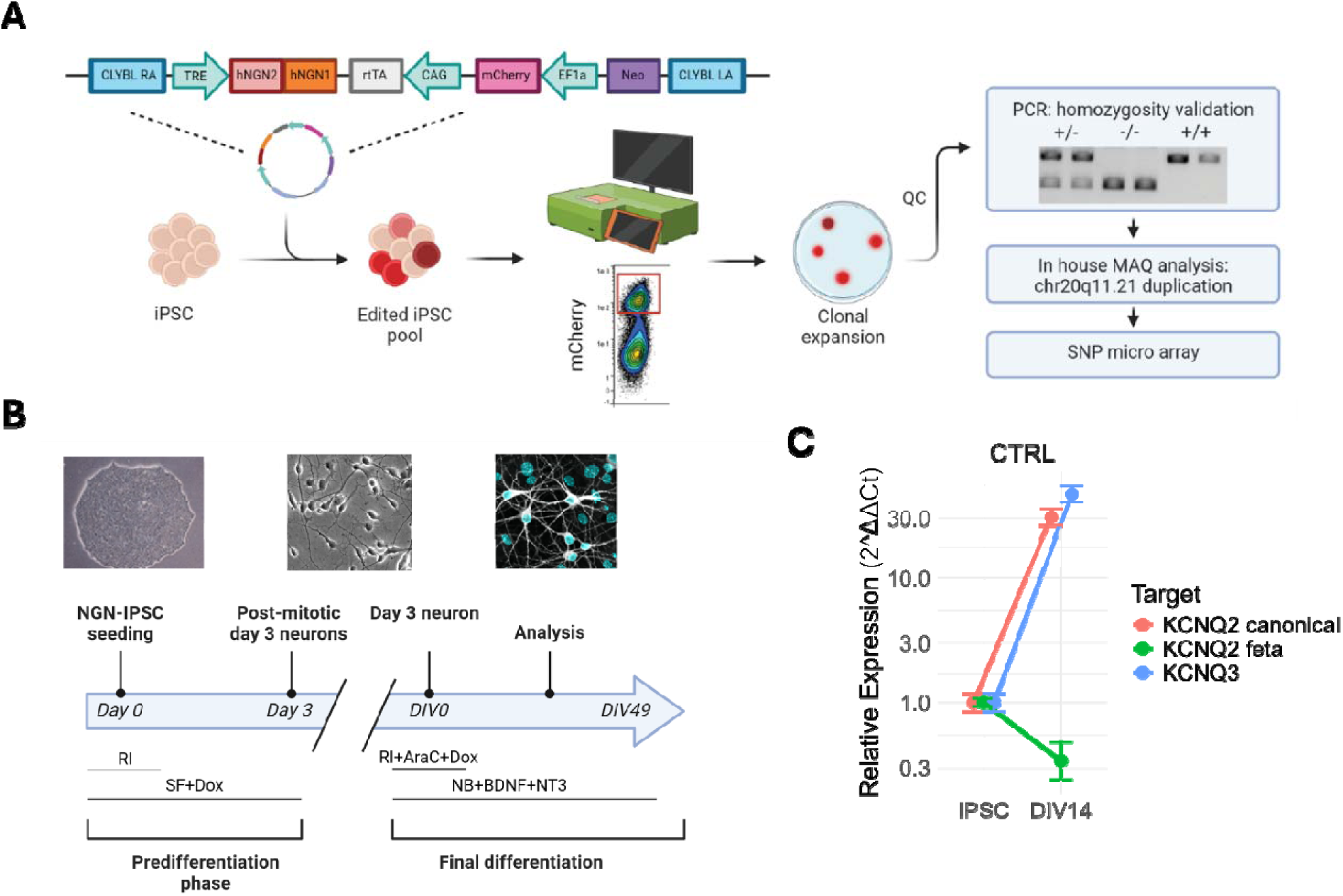
Generation of and characterization of KCNQ2-LOF iNeurons. A) Graphical overview of stable NGN-iPSC generation and quality control. B) Protocol for iNeuron generation, showing NGN-iPSC, day 3 neurons, and MAP2 positive DIV14 iNeurons. RI: Rock inhibitor, DF: Stem Flex, Dox: Doxycycline, AraC: cytosine arabinoside, NB: Neurobasal, BDNF: Brain derived neurotrophic factor, NT3: neurotrophin-3. C) RT-qPCR analysis of fetal KCNQ2, canonical KCNQ2, and KCNQ3 transcripts in iPSC and DIV14 iNeurons of pooled controls (WT and ISO_G290D). Values are normalized to respective target in iPSC lysate. N=8/4

The expression of the short fetal and canonical *KCNQ2* transcripts (NM_172109 and NM_172107, respectively) and *KCNQ3* transcript (NM_004519) were validated in iPSCs and day *in vitro* (DIV) 14 neurons of the control lines (WT and ISO_G290D). In line previous reports, the short fetal KCNQ2 transcript showed a decreasing expression pattern during neuronal maturation, while the canonical *KCNQ2* and the *KCNQ3* transcripts showed an increase in expression levels (Fig. 1B, right) (15, 23).

### The KCNQ2-DEE G290D variant increases neuronal excitability under stimulation at the single-cell level

To investigate whether severe Kv7.2 LOF affects electrophysiological properties in iNeurons, we performed whole-cell patch-clamp recordings using the isogenic pair in our cohort (KCNQ2-DEE G290D and ISO_G290D), to minimize confounding effects from genetic background, between DIV 18–28. Voltage-clamp recordings showed no differences in sodium or potassium current density between lines (Fig. 2A-C). Current-clamp recordings showed that both lines were electrically active, capable of generating spontaneous action potentials (APs) at similar rates (~70% of neurons, with firing frequencies of 2.1 ± 0.5 APs/s for ISO_G290D and 2.2 ± 0.4 APs/s for G290D; Fig. 2D-E), with no significant differences in passive membrane properties (Fig2, Table 2). However, upon slow depolarizing current ramps from −20 to +200 pA to assess neuronal excitability, KCNQ2-DEE G290D iNeurons fired significantly more APs than ISO_G290D (p < 0.05; Fig. 2G–H). Single AP properties showed no differences in the AP threshold, rheobase, amplitude, width, and fast afterhyperpolarization (fAHP) (Table 2). These results indicate KCNQ2-DEE G290D iNeurons display increased excitability when stimulated.

**Fig. 2.**
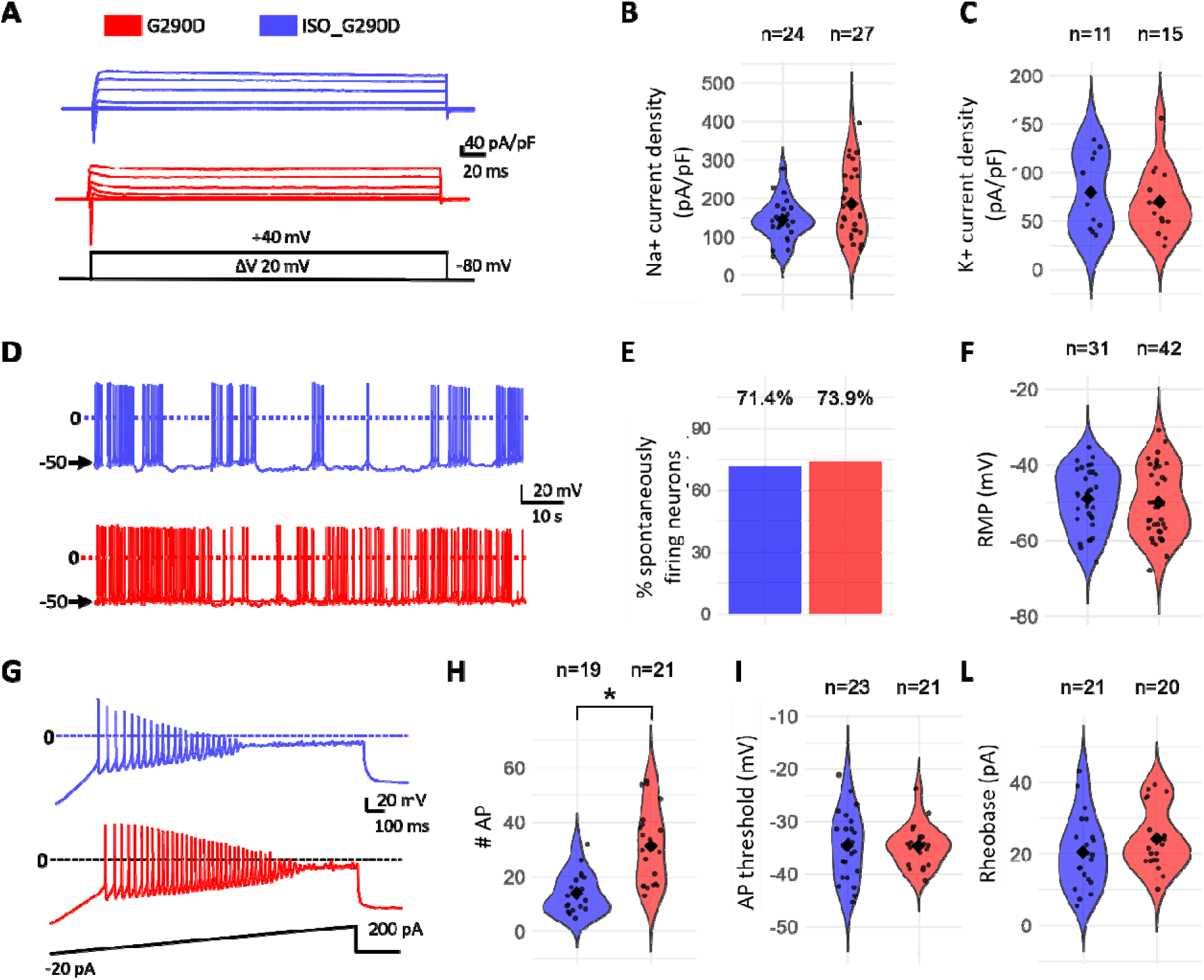
Electrophysiological properties of KCNQ2-DEE G290D and ISO_G290D iNeurons. A) Representative current traces of KCNQ2-DEE G290D iNeurons (red) and ISO_G290D iNeurons (blue) evoked by the indicated voltage protocol. B) Peak sodium (Na^+^) current density calculated at −10 mV and (C) potassium (K^+^) current density calculated at +20mV. D) Representative traces of iNeurons spontaneous activity recorded in whole-cell current clamp. E) Percent of spontaneously firing neurons and (F) resting membrane potential (RMP) G) Representative current-clamp recordings during ramp depolarization. H) Number of action potentials (APs), (I) AP threshold and (L) rheobase. In panels B, C, F, H, I and L each bar shows the mean ± SEM from the indicated number of cells recorded in three rounds of iNeuron differentiation (N=3). *p<0.05.

### KCNQ2-LOF iNeurons show earlier onset of firing and increased excitability during early maturation

To investigate the electrophysiological effects of *KCNQ2*-LOF variants on a population-wide level over time, we used high-density multielectrode arrays (HD-MEA). Neuronal cultures were recorded two to three times per week from DIV10 until DIV49. All genotypes initiated spontaneous firing between DIV10 and DIV13, with mean firing rate (MFR) and active electrode area (AA, percentage of electrodes recording spikes) significantly increasing over time (p<0.001, and net positive first derivatives), indicating electrophysiological maturation (Fig. S2 and Fig. S3A,B). Next, we examined whether Kv7.2 LOF affected the timing of spontaneous firing onset. During early maturation (DIV10–24), KCNQ2-LOF cultures displayed a significantly higher AA than the pooled WT and ISO_G290D (p < 0.001, except A294V; p = 0.39), suggesting that LOF neurons become electrophysiologically active earlier during maturation.

Given that all iNeuronal cultures displayed active firing by DIV13, we restricted our following analysis to recordings from this timepoint onward to ensure inclusion of active neurons across all genotypes. Throughout maturation, all genotypes demonstrated dynamic changes in electrophysiological behavior, as illustrated in the smooth functions shown in Fig. S2. Given the complexity and non-linearity of the neuronal activity changes, generalized additive models (GAM) provided a flexible and robust analytical framework to accurately characterize these effects (24). We performed pairwise comparisons between the *KCNQ2*-LOF lines and the pooled control, and generated comparative genotype-specific significance and fold change heatmaps for all parameters over time to visualize these dynamical changes (see Fig. S4 for GAM pipeline).

Comparative heatmaps revealed consistent temporal trends in neuronal activity across all KCNQ2-LOF mutant lines (Fig. 3B-C). To identify shared patterns among *KCNQ2*-LOF variants, parameters were selected based on two criteria; (i) a significant difference when comparing pooled KCNQ2-LOF vs. pooled control iNeurons (Fig. 3B), and (ii) a consistent directionality of the effect in at least three individual genotypes compared to the pooled control (Fig. 3C). All KCNQ2-LOF lines exhibited a significant increase in MFR and FR variability (coefficient of variance (CV)), particularly during early maturation, with no consistent differences in spike amplitude (Fig. 3C). Together with the patch clamp data, these results support a hyperexcitable phenotype in *KCNQ2*-LOF neurons during early maturation.

**Fig. 3.**
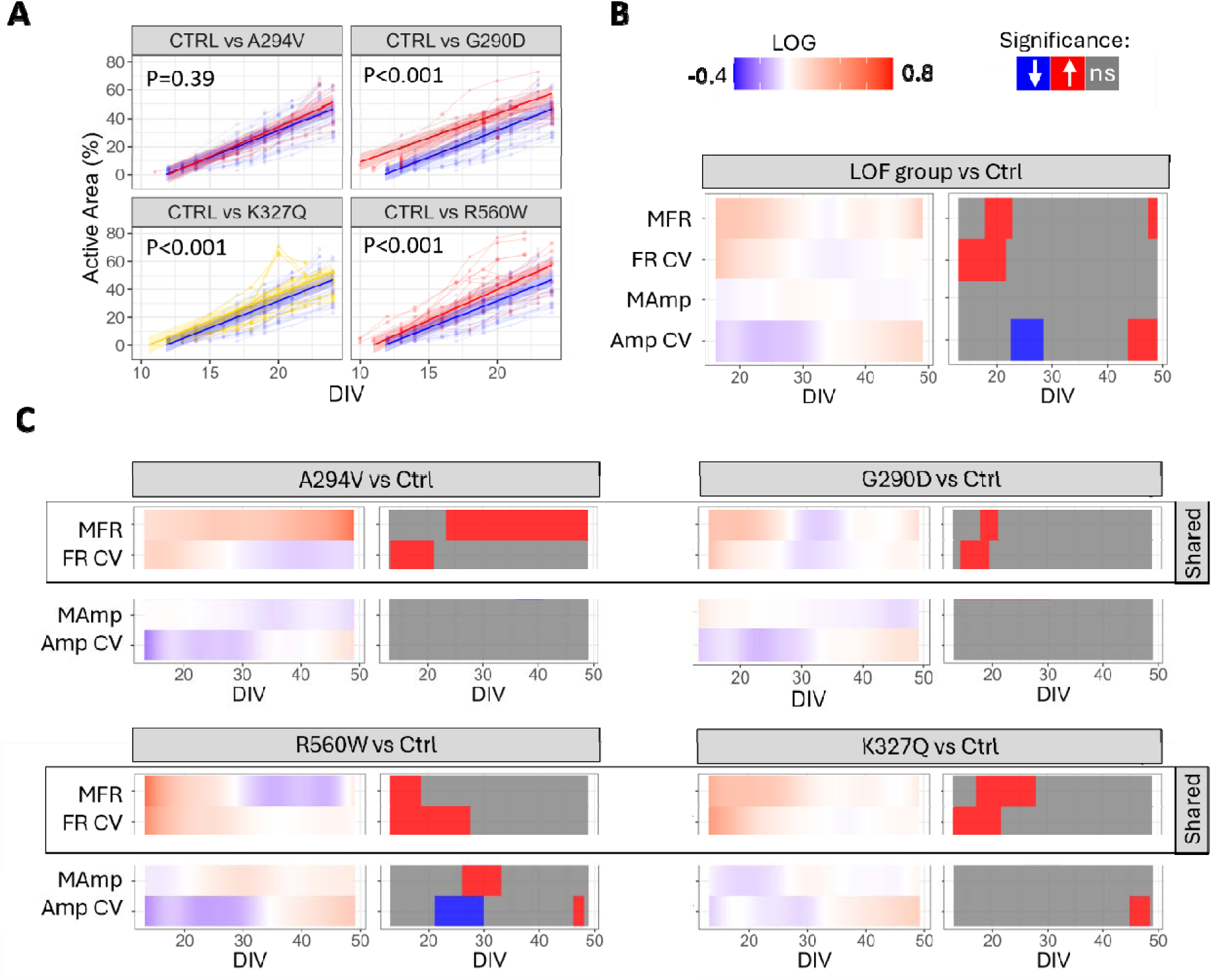
Longitudinal comparison of HD-MEA activity parameters of KCNQ2-LOF iNeurons to its control. A) Longitudinal modeling of active area percentage from DIV10–24. Thin lines show individual trajectories; solid lines show mixed-effects model predictions ±95% CI. Genotype vs. control (CTRL) comparisons used estimated marginal means with Dunnett’s adjustment. Blue: CTRL; Red: KCNQ2-DEE; Yellow: KCNQ2-SeLFNE. B–C) Comparative Heatmaps show fold-change (FC) vs. CTRL; pairwise comparisons indicate significance. B) CTRL versus Grouped LOF data; C) CTRL versus Genotype-specifi data. Positive FC and pairwise values (red) = increase; negative (blue) = decrease. n=39/11 CTRL, 16/6 K327Q, 13/6 A294V, 19/5 G290D, 19/7 R560W. MFR: mean firing rate; FR CV: firing rate coefficient of variance; MAmp: mean amplitude; Amp CV: amplitude coefficient of variance.

### KCNQ2-LOF iNeurons exhibit shared stable and biphasic network behavior patterns

HD-MEA network analysis revealed that all cultures developed synchronized network bursts between DIV14–21, with burst frequency (BF) increasing over time (p<0.001, and net positive first derivatives), reflecting neuronal maturation (25) (Fig. S3C and S5). As described by Wagenaar DA et al., our neuronal cultures also exhibited an evolution in burst wave form and burst patterns throughout development (26). Burst waveforms evolved from synchronous single-peak network bursts to complex synchronous reverberating bursts, characterized by sequential mini-bursts occurring within a single network burst (Fig. S6). Early in maturation, long-tailed network bursts were observed in ISO_G290D, G290D and R560W (Fig. S7). After DIV28, long-tailed bursts were no longer observed. To accurately quantify network parameters influenced by these burst tails, a 95% peak-to-baseline cut-off was applied for data up to DIV28, in addition to the standard 80% cut-off used throughout the study (Table 3 and Fig. S7). Due to the non-normal distribution of ISI outside of bursts, we analyzed the median as well as the 10th (P10) and 90th (P90) percentiles instead of the mean.

Using the same inclusion criteria to define shared patterns, we identified nine parameters consistently altered in *KCNQ2*-LOF lines compared to controls (Fig. 4A-B, Fig. S8). While some network parameters showed a consistent increase or decrease throughout maturation, others displayed a dynamic biphasic pattern with opposing trends. Dynamic parameters seemed to transition around the same DIV within each genotype, allowing for the identification of two distinct phases (further named phase 1 and 2) in network behavior (Fig. 4B). Throughout most of the recording period, all *KCNQ2*-LOF cultures exhibited shorter inter-burst intervals (IBI), higher BF, a higher percentage of spikes occurring within bursts, greater variability in spike timing within bursts (burst ISI CV), and a shorter ISI P90 outside of bursts (non-burst ISI). During phase 1, these changes were accompanied by longer bursts (BD) that contained a larger number of spikes (SPB), with longer ISI (burst_ISI) and greater ISI variability outside of the bursts (non-burst ISI CV), which shifted in the opposite direction during phase 2 (Fig. 4, Fig. S6). The transition between phases varied by line, with the SeLFNE variant K327Q transitioning almost immediately (~DIV20), followed by the DEE variants R560W (~DIV30), G290D (~DIV40) and A294V (~DIV49) (Fig. 4B).

**Fig. 4.**
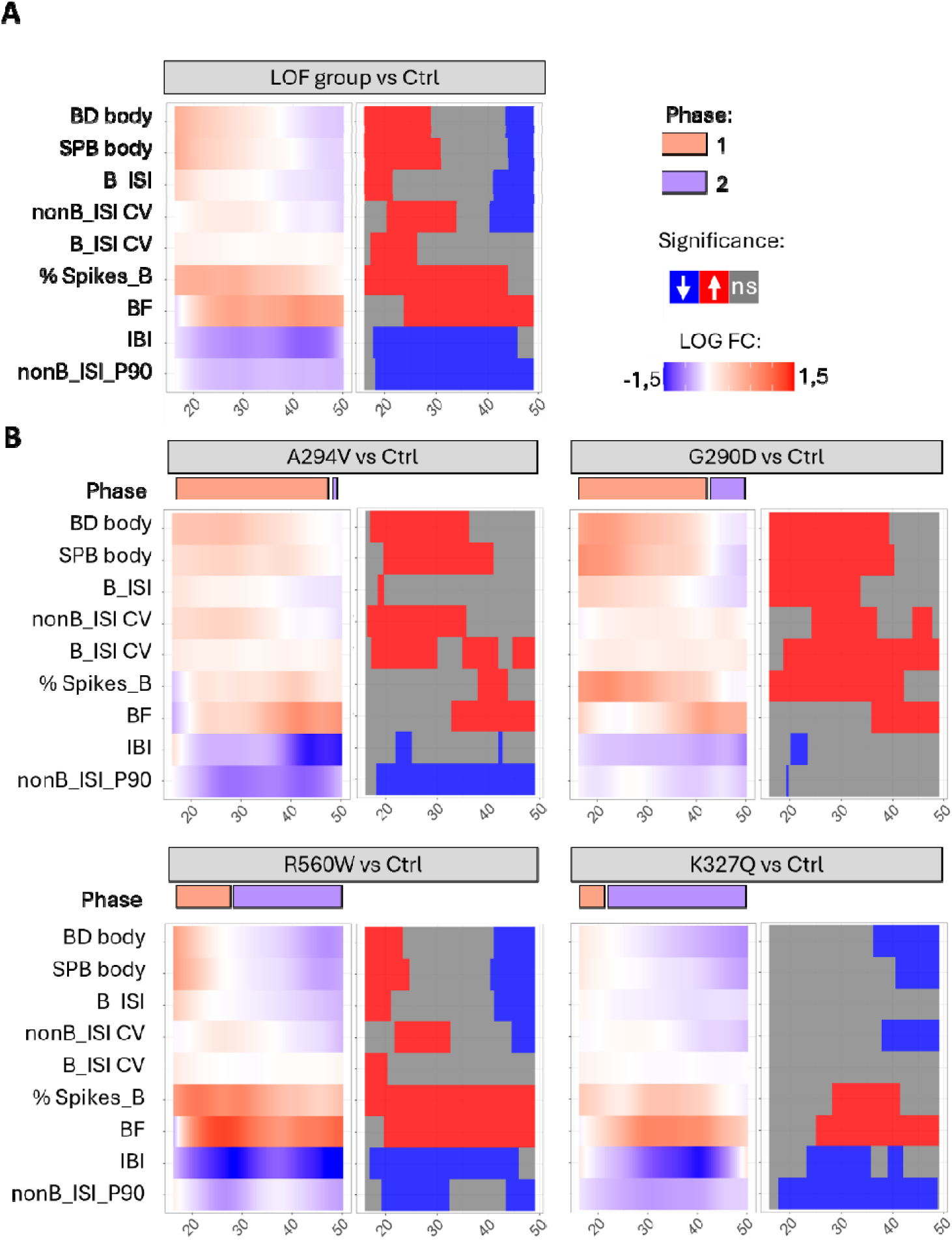
Longitudinal comparison of shared HD-MEA network parameters of KCNQ2-LOF iNeurons to its control. A) Grouped LOF data. B) Genotype-specific data. Comparative Heatmaps show fold-change (FC) vs. CTRL; pairwise comparisons indicate significance. Positive FC and pairwise values (red) = increase; negative (blue) = decrease. n=39/11 CTRL, 16/6 K327Q, 13/6 A294V, 19/5 G290D, 19/7 R560W. B: burst, BD: Burst duration, SPB: spikes per burst, ISI: interspike interval, nonB_ISI: burst inter spike interval outside of burst, CV: coefficient of variance, IBI: inter burst interval, BF: burst frequency, % spikes_B: percentage of spikes within bursts.

### RTG differentially affects mutant-relevant parameters in early and late phases of neuronal maturation

Given the consistent and biphasic network pattern observed in *KCNQ2*-LOF iNeurons, we used the Kv7 channel activator retigabine (RTG) to determine which mutant-associated parameters at phase 1 and 2 resulted directly from reduced M-current and which from adaptive mechanisms. First, we validated our model by showing that a single 10 µM RTG exposure at DIV 21 inhibited over 80% of spontaneous firing in the KCNQ2-G290D iNeurons using whole-cell patch-clamp (Fig. 5A). Second, we performed a concentration-response (CR) at both phases using HD-MEA, to select a RTG concentration that affected AA and burst parameters without completely silencing network bursting (all recorded wells still had to show synchronized network burst activity). The AA-CR curve confirmed the silencing effectiveness of 10 µM RTG at phase 1 and phase 2 (Fig. 5B). Importantly, neuronal silencing by the highest RTG concentration was reversible by application of 20 µM XE991, a Kv7 channel blocker (Fig. 5B), demonstrating that RTG-induced silencing was not due to cell death but rather to functional modulation of Kv7 channels. Interestingly, we found a nearly two-fold reduction in RTG potency during phase 2 (IC50 = 849 nM) compared to phase 1 (IC50 = 473 nM) (Fig. 5B). From the CR-curves, a concentration of 100 nM for phase 1 and 300 nM for phase 2 was selected, both approximating the IC20.

**Fig. 5.**
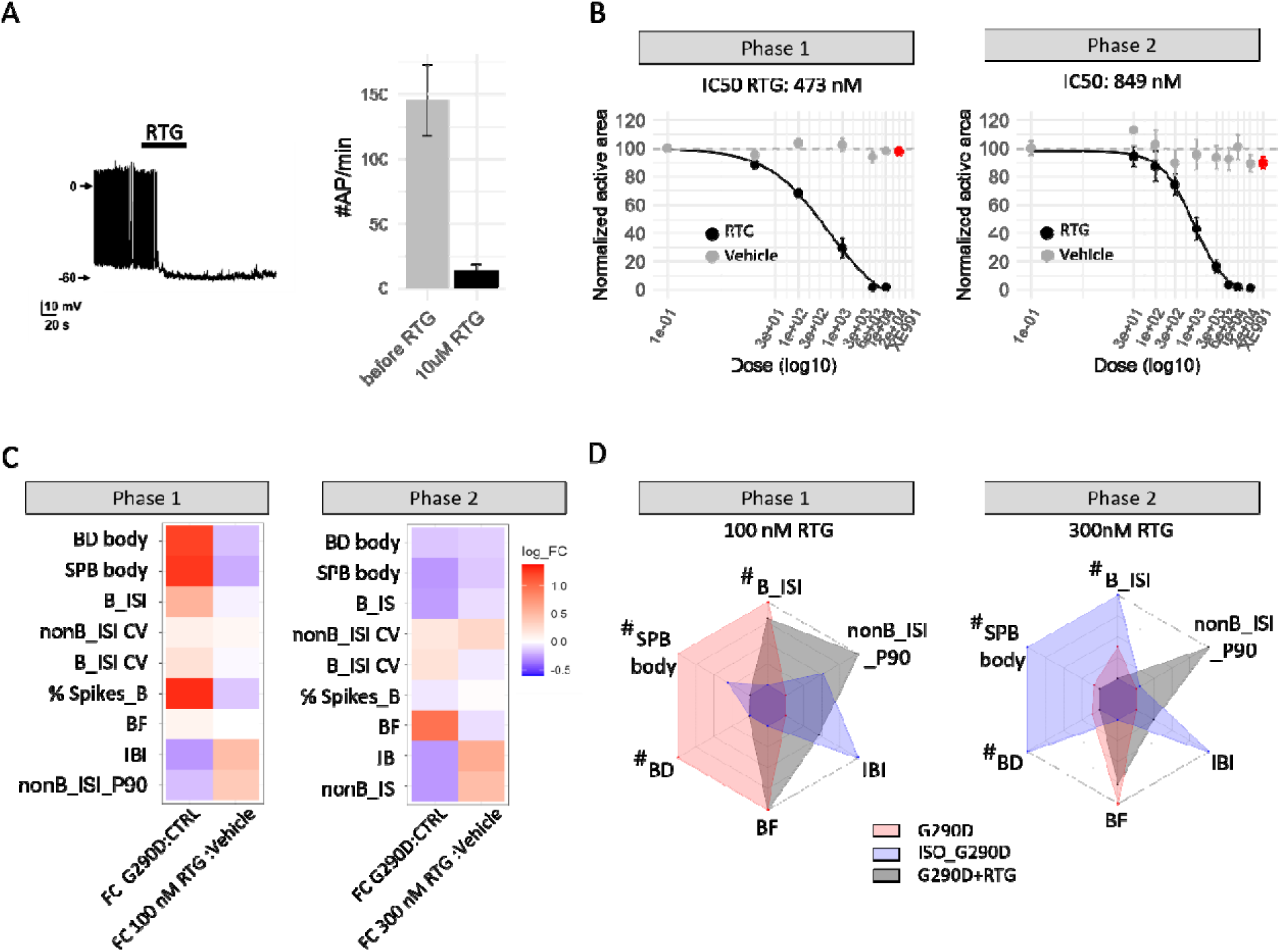
Effect of RTG treatment on G290D during phase 1 and phase 2. A) Left, Representative spontaneous firing from whole-cell current clamp of G290D iNeurons exposure to 10 µM retigabine (RTG) at indicated bar at DIV21. Right, barplot of mean AP per minute before and after 10 µM RTG treatment. B) IC50 curves for RTG on G290D and XE991 (red) recovery during phase 1 and phase 2. N=9/3 for RTG, N=7/3 for Vehicle. C) Fold change heatmap of HD-MEA network parameters comparing untreated G290D with ISO_G290D in the first column, and the RTG-induced changes to the G290D profile in the second column during phase 1 (Left) and 2 (Right). D) Radar maps of HD-MEA network parameters showing G290D (pink), ISO_G290D (blue) and G290D treated with RTG (grey) at phase 1 (left) and phase 2 (right). N=9/3 for RTG, N=7/3 for Vehicle. # indicates biphasic parameters, * indicate phase independent parameters.

During phase 1, RTG but not vehicle induced network changes in G290D that pushed several phase 1 mutant parameters towards ISO_G290D values, notably also reducing biphasic parameters BD, SPB and burst ISI (Fig. 5C-D). Additionally, RTG reduced the severity of phase-independent mutant parameters non-burst ISI P90 and IBI. These results are indicative of Kv7.2 LOF playing a direct role in the phase 1 phenotype. In contrast, during phase 2, G290D exposure to 300 nM RTG induced similar overall effects on measured parameters compared to phase 1, resulting in the rescue of the phase independent network parameters, but the exacerbation of the biphasic network parameters (Fig. 5C-D).

These results indicate that, while decreased Kv7.2 function primarily drives the electrophysiological phenotypes in phase 1, the characteristics of more mature mutant networks are partly the result of a (dysfunctional) homeostatic response to the network imbalances, rather than being solely dictated by Kv7.2 dysfunction.

### KCNQ2-DEE G290D Neurons Exhibit Phase-Specific Transcriptomic Dysregulation and Kv7 Channel Expression

To investigate transcriptomic changes underlying the biphasic network behavior, we performed RNA-seq analysis at phase 1 (DIV21) and phase 2 (DIV49) comparing KCNQ2-G290D to ISO_G290D. PCA showed clear genotype-driven separation (PC2) at both timepoints with greater divergence at DIV21 (Fig. 6A), further supported by the nearly twice as many differentially expressed genes at DIV21 compared to DIV49 (1365 vs. 723; FDRl1<l10.05 and |log2FC|>0.58) (Fig. S9A). Furthermore, epilepsy-associated genes were enriched at both timepoints, while cardiac disease-related gene enrichment was absent, supporting the specificity of transcriptomic changes to epilepsy-relevant pathways (Fig. S9B).

**Fig 6.**
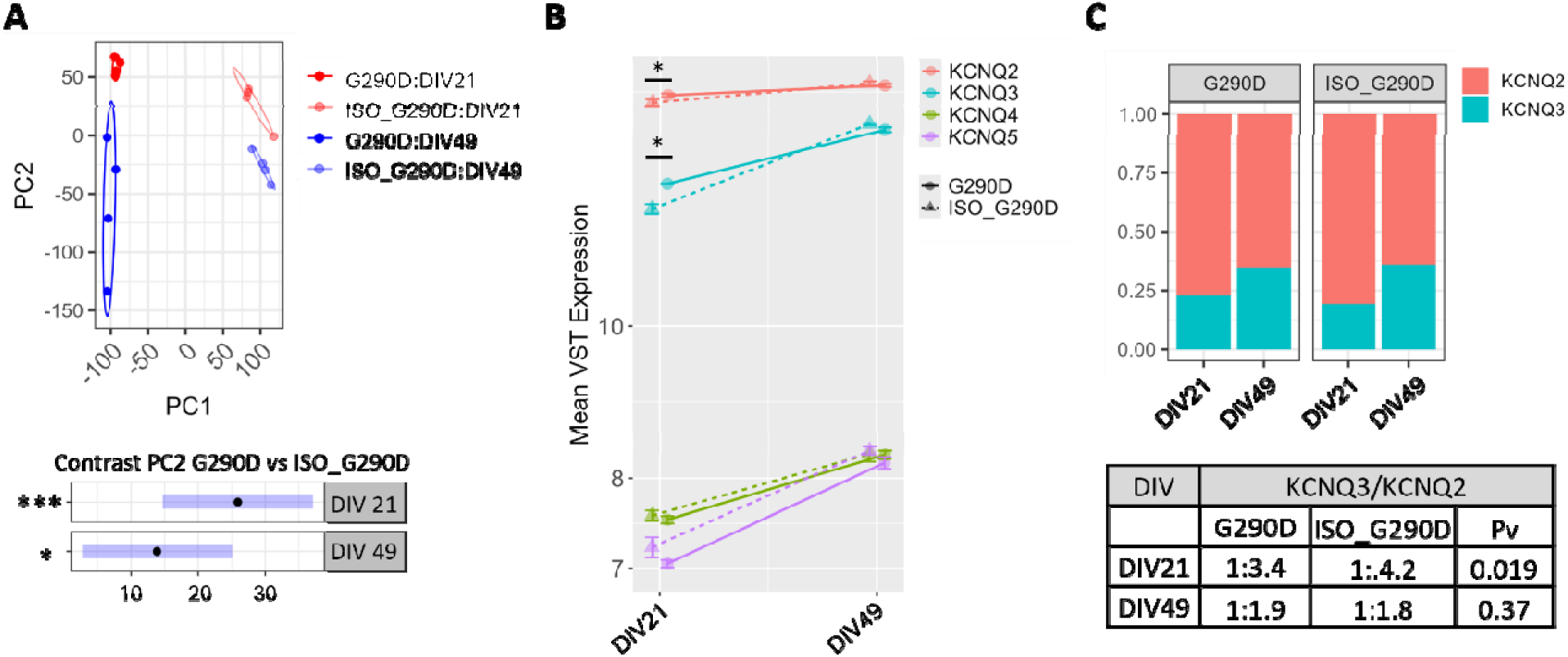
Transcriptomic analysis of KCNQ expression in KCNQ2-G290D and ISO_G290D iNeurons in early and late maturation. (A, top) Principal component analysis (PCA) plot of variance-stabilized expression (VST) values of all genes with non-zero variance across samples, showing separation of samples by genotype along PC2 and by time along PC1. (A, bottom) PC2 contrast calculation for DIV21 and DIV49. *p<0.05, ***p<0.001. (B) Line plots showing the mean VST expression levels + SEM of KCNQ2, KCNQ3, KCNQ4, and KCNQ5 genes over time in G290D and ISO_G290D iNeurons. (C) Proportion plots illustrating the ratio of KCNQ3 to KCNQ2 expression in G290D and ISO_G290D iNeurons at DIV21 and DIV49. The accompanying table summarizes the mean ratios and statistical comparisons between genotypes at each timepoint.

Next, we evaluated KCNQ expression in the RNA sequencing data. Both genotypes showed increasing *KCNQ2*-5 expression over time (Fig. 6B). Notably, the KCNQ2:KCNQ3 expression ratio was highest at phase 1 for both genotypes and decreased substantially in phase 2 (Fig. 6C). Among the 20 highest expressed potassium channel genes, KCNQ2 was the most highly expressed gene during both phases, with *KCNQ3* ranking second or third (Table 4). Notably, *KCNQ2* and *KCNQ3* expression was significant upregulated in G290D iNeurons at phase 1, but not phase 2 (Fig. 6B), accompanied by a shift in the KCNQ3/KCNQ2 expression ratio, from 1:4.2 (ISO_G290D) to 1:3.4 (G290D, p=0.019) (Fig. 6C). Given that Kv7.2/Kv7.3 hetero-tetramers generate an M-current density approximately 3.8 times greater than homomeric channels, these changes in gene expression likely represent an adaptive response compensating for the loss of functional M-current in G290D iNeurons.

### KCNQ2-DEE G290D ineurons Display a Compensatory Shift in Potassium Channel Gene Expression during Early Maturation to Maintain Potassium Homeostasis

To explore the broader biological processes affected in developing KCNQ2-DEE G290D iNeurons, we performed gene ontology (GO) enrichment analysis on genes with significant time × genotype interaction (FDR<0.05). Enriched processes included potassium ion transport, neuron apoptotic process, dopamine transport, calcium ion import across the plasma membrane, and retrograde axonal transport (Fig. 7A). The top GO hit ‘potassium ion transport’ was predominantly driven by differential expression of potassium channel subunits genes and their modulators (Fig. 7B). These changes were most pronounced at phase 1, reducing or normalizing by phase 2. Notably, we observed upregulation of several genes encoding for subunits of potassium efflux channels or modulators enhancing efflux, along with downregulation of genes encoding for subunits of potassium influx channels or channel modulators reducing potassium efflux. This suggests an adaptive cellular response aimed at compensating for the loss of the Kv7.2 M-current (potassium efflux) to maintain potassium homeostasis.

**Fig 7.**
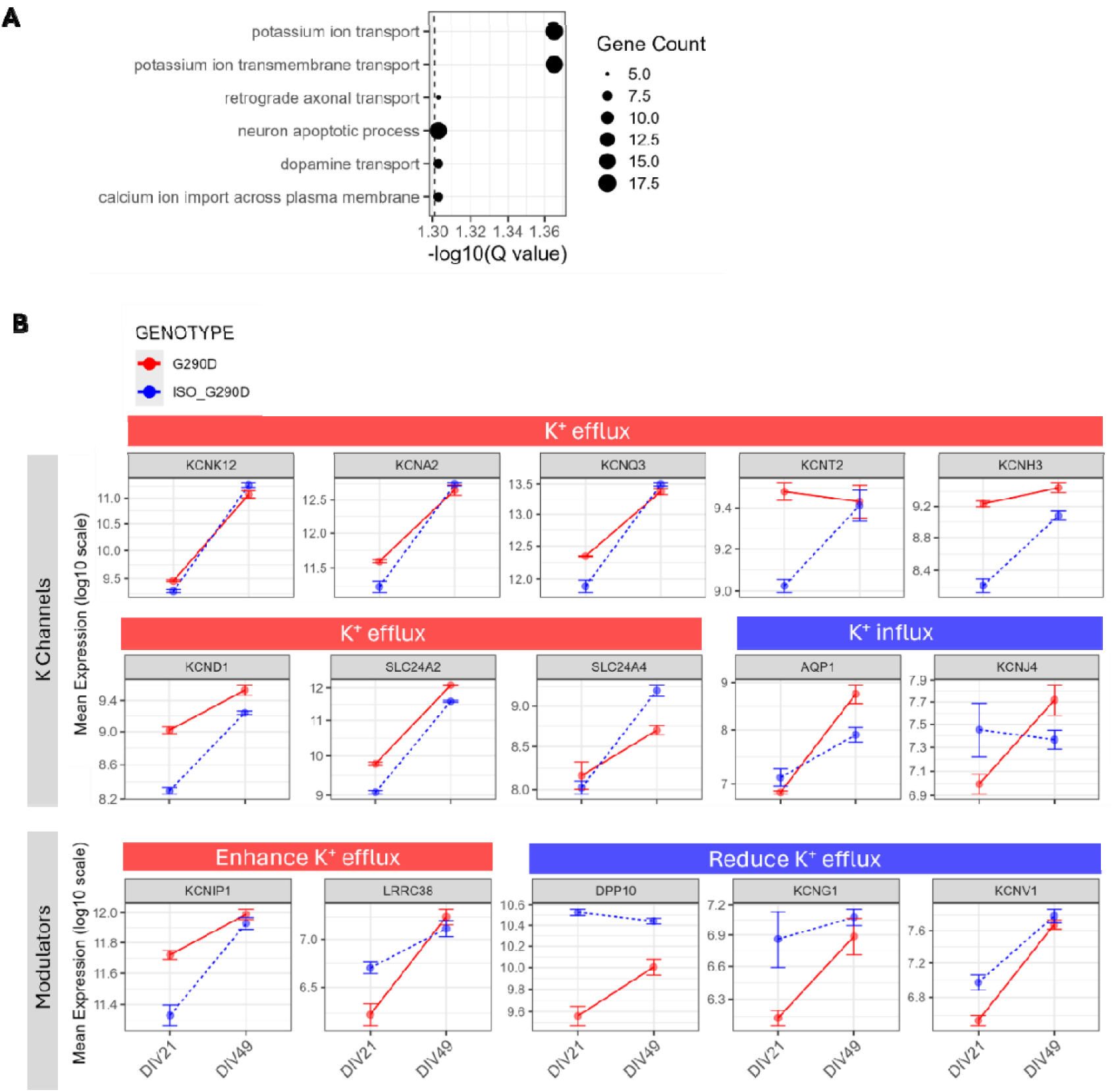
Transcriptional dysregulation of potassium channel gene expression in KCNQ2-DEE G290D iNeurons at early and late maturation. (A) Gene ontology (GO) enrichment analysis of significant genes, illustrating the enriched biological processes, with a particular emphasis on potassium ion transport. (B) Line plots showing the normalized temporal expression profiles of the genes included in the potassium ion transport GO term across developmental phases. Genes encoding subunits of potassium channels are separated into subunits that are part of efflux (red) or influx (blue) potassium channels. Genes encoding for potassium channel modulators are separated into modulators that enhance (red) or reduce (blue) the efflux.

### KCNQ2-DEE G290D iNeurons display phase dependent synaptic gene expression signatures aligned with network activity changes

We next performed GO enrichment analysis of biological processes at each timepoint. The top 10 biologically relevant GO terms were almost completely shared between timepoints and were dominated by synaptic processes (Fig. S10A). Synapse specific GO term analysis (SynGO) of up- and downregulated DEGs separately for each timepoint (FDR<0.05 and |log2FC|>0.58) revealed a phase-specific pattern: at DIV21, synaptic enrichment was driven mainly by upregulated genes, while at DIV49 it was driven by downregulated genes (Fig. S10B). Of the 49 synapse-related DEGs shared across phases, only two switched direction of expression, indicating that distinct gene sets underlie the observed phase-specific signatures (Fig. S10C).

To better understand the functional implications of these uniquely enriched up and down regulated DEGs, we reviewed their reported roles in glutamatergic synaptic function (Table 5). Functional impact on synaptic processes of biologically relevant DEGs is schematically summarized in Fig. 8A. In phase 1, upregulated synaptic DEGs pointed to enhanced glutamate release dynamics and postsynaptic receptor modulation, with overall reinforcement of excitatory synaptic stability and strength. Presynaptically, the transcriptomic profile indicated enhanced recruitment, mobilization, and fusion of synaptic vesicles, together with upregulation of calcium and cAMP signaling pathways, suggesting a presynaptic state primed for elevated glutamatergic release. Postsynaptically, upregulation of several modulatory factors enhancing NMDA and AMPA receptor signaling was observed. Since enhanced NMDA and AMPA signaling is known to increase burst duration and burst frequency, respectively, we examined the receptors’ gene expression in the transcriptome data. This revealed a downregulation of GRIA2-4 and GRIN2A, GRIN2B and GRIN3B genes, possibly reflecting a compensatory mechanism aimed at dampening excessive excitatory signaling (Fig. 8B). These opposing transcriptional changes complicate predictions regarding the net functional effect on NMDA and AMPA signaling.

**Fig 8.**
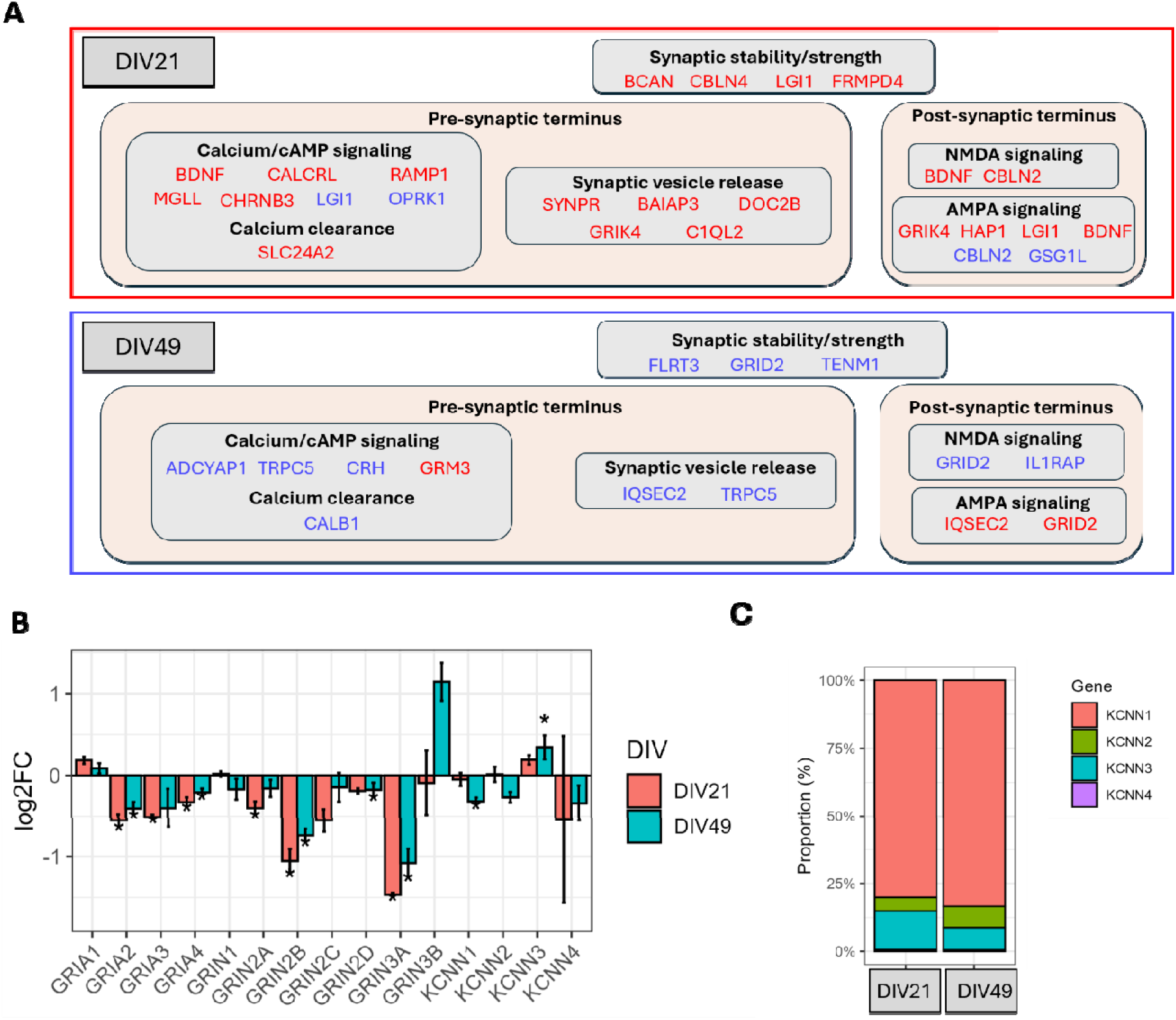
Transcriptional dysregulation of synaptic gene expression in KCNQ2-DEE G290D iNeurons in early and late maturation. A) Schematic representation of the biologically relevant uniquely upregulated synaptic DEGs at DIV21 (red box), and uniquely downregulated synaptic DEGs at DIV49 (blue box) identified by SynGO term enrichment analysis and their putative roles in glutamatergic synaptic function. DEGs are categorized according to their functional involvement in synaptic processes. DEGs highlighted in blue are associated with a decrease in the assigned synaptic process, while DEGs in pink are associated with an increase. B) Bar plot shows the log2 fold change in expression for AMPA/NMDA receptor subunits genes (GRIA1-4 and GRIN1-3) and SK channel genes (KCNN1–4) in G290D mutant neurons, compared to ISO_G290D, at DIV21 (pink) and DIV49 (blue). Data is presented as mean + SEM, *FDR<0.05. C) Proportion plots illustrating the ratio of KCNN1-4 expression in ISO_G290D neurons at DIV21 and DIV49.

Phase 2-specific downregulated synapse-related DEGs pointed towards a decreased glutamate release and postsynaptic receptor modulation (Fig. 8A). Downregulation of genes associated with synaptic adhesion and scaffolding suggested weakening of synaptic structure and connectivity. Presynaptically, downregulated genes suggested a diminished calcium/cAMP signaling and synaptic vesicle release, indicating a reduced glutamate release. Postsynaptically, a similar decrease of NMDA receptor genes was observed as in phase 1, however now accompanied by downregulation of positively modulating genes, suggesting decreased NMDA signaling (Fig. 8A-B), consistent with the observed shortening of BD.

The marked increased BF characteristic of phase 2 could reflect enhanced AMPA receptor signaling. However, the transcriptomic profile was complex, with decreased AMPA receptor gene expression alongside increased genes promoting AMPA receptor surface expression. To further explore other postsynaptic mechanisms, we assessed small conductance calcium-activated potassium (SK) channel genes (*KCNN1-3*) as reduced SK function is known to elevate burst frequency and decrease ISI (Fig. 8B-C). KCNN1 was significantly downregulated, whereas KCNN3 was upregulated. As KCNN1 accounts for approximately 83% of total KCNN expression at DIV49, and KCNN3 only for 8%, a net KCNN reduction potentially contributes to the increased burst frequency and shortened ISI observed at phase 2 (Fig. 8C).

Together, these findings highlight a dynamic, biphasic regulation of synaptic gene expression in G290D iNeurons, with early maturation marked by enrichment of genes promoting glutamate release, and late maturation characterized by enrichment of genes associated with reduced glutamate release.

### KCNQ2-LOF iNeurons show a greater reduction in presynaptic density over time

Given the opposing trends in electrophysiological network activity and glutamatergic signaling suggested from the HD-MEA and transcriptome data across maturation phases, we next assessed whether these were also reflected at the synaptic structure. We quantified total pre-synaptic, post-synaptic, and overall synaptic density (colocalization of Bassoon and Homer1) with MAP2 as a neurite marker (Fig. 9A).

**Fig 9.**
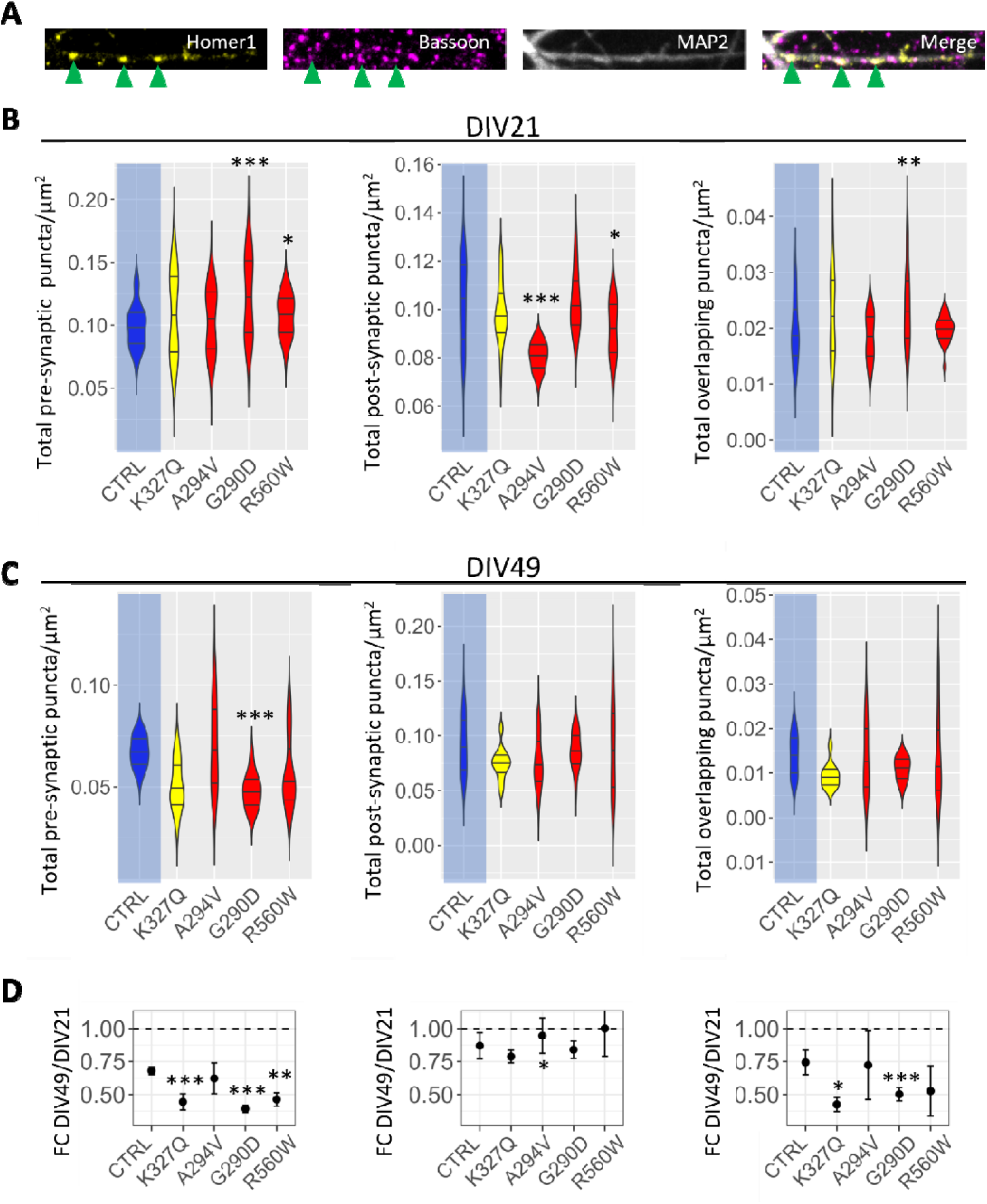
Quantification of synaptic density across KCNQ2-LOF iNeurons at DIV21 and DIV49. A) Representative images of dendrites at DIV21 MAP2 (dendrites, grey), Homer1 (post-synaps, Yellow) and Bassoon (pre-synaps, magenta). Green arrowheads = co-localized puncta. B, C) Quantification of total pre-, post-, and synaptic density at DIV21 (B) and DIV49 (C). Control (CTRL) in blue. D) Fold change in pre-synaptic density (DIV49:DIV21 ratio). Data represented as mean ± SEM. Statistical analysis was performed using the interaction linear mixed models, genotype x DIV; post hoc = Benjamini-Hochberg. * p_adj < 0.05 vs. CTRL.

At DIV21, KCNQ2-G290D and KCNQ2-R560W iNeurons showed a significant increase in pre-synaptic density (p<0.001 and p<0.05, respectively), whereas KCNQ2-A294V and KCNQ2-R560W showed reduced post-synaptic density (p <0.001 and p<0.05, respectively) (Fig. 9B). Only KCNQ2-G290D demonstrated an increase in total synaptic density (p<0.01). By DIV49, a significant decrease in pre-synaptic density was observed in KCNQ2-G290D alone (p<0.001) (Fig. 9C). However, when comparing synaptic changes over time (DIV49:DIV21), all KCNQ2-LOF lines, except A294V, displayed a significantly greater reduction in pre-synaptic density over time (K327Q and G290D: p<0.001, R560W: p<0.01) (Fig. 9D). The absence of a significant decline in KCNQ2-A294V may reflect its delayed and still ongoing transition to phase 2.

### KCNQ2-LOF variants cause alterations in AIS Dynamics Under Naïve and Chronic Depolarizing Conditions

Kv7.2 channels interact with Ankyrin-G and are enriched at the AIS. In response to prolonged changes in neuronal excitability, the AIS undergoes homeostatic structural plasticity, adjusting its length and position to stabilize neuronal firing. As chronic M-current suppression in primary hippocampal neurons has been shown to induce AIS structural adaptations (27), we examined whether *KCNQ2*-LOF variants affect AIS length or location, using Ankyrin-G and MAP2 as markers for AIS and cell soma identification, respectively. Additionally, we assessed if AIS plasticity was affected in *KCNQ2*-LOF iNeurons by the established method of inducing chronic depolarization via KCl treatment at 10mM for 48 hours (Fig. 10A) (28–31).

**Fig 10.**
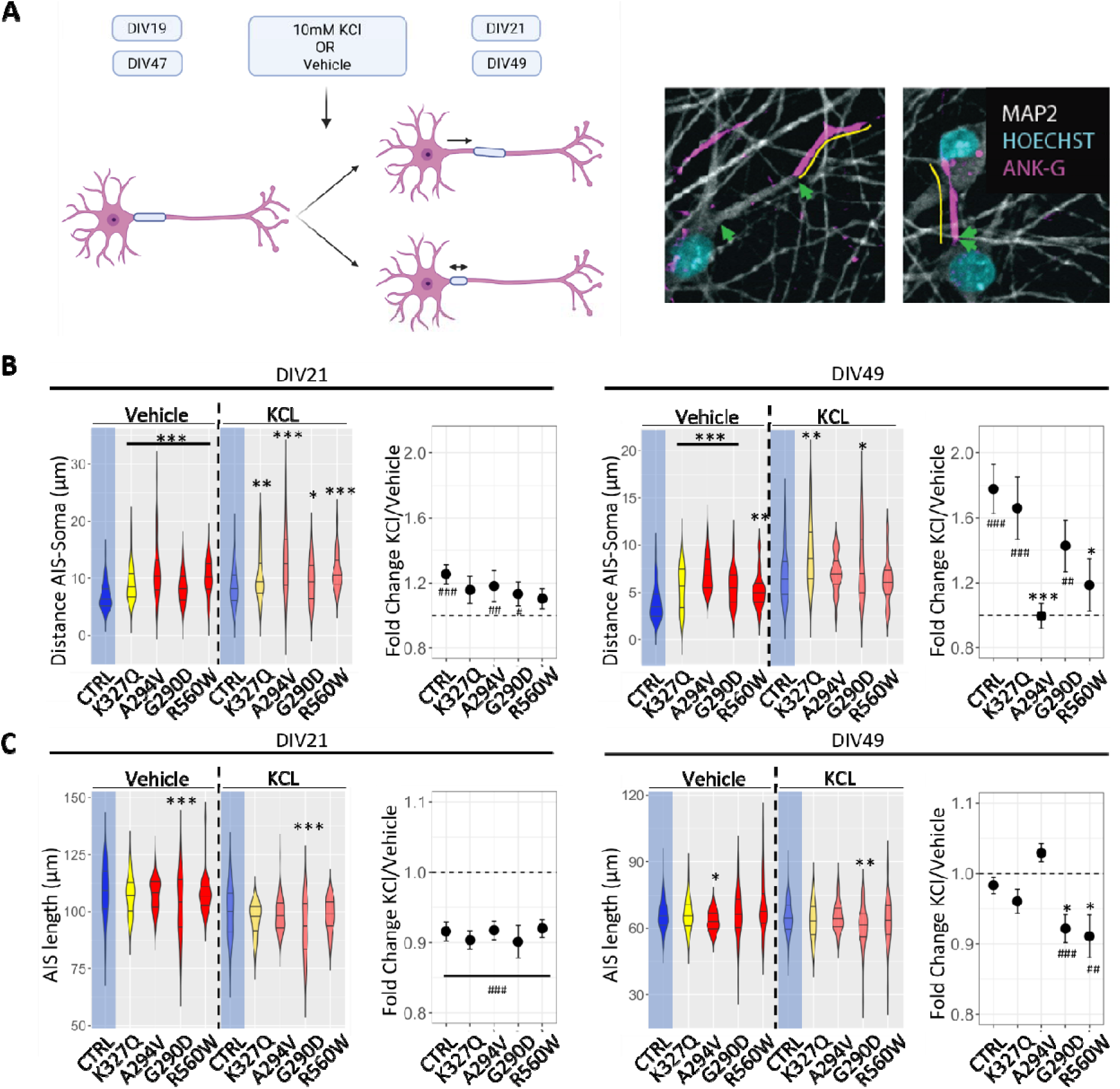
Structural AIS modifications and plasticity in KCNQ2-LOF iNeurons. A) Experimental overview: DIV19 and DIV47 neurons treated with 10mM KCl or vehicle for 48h; stained for AIS (AnkG, magenta), dendrites (MAP2, grey), nuclei (DAPI, cyan). Yellow line: AIS length; green arrows: soma-to-AIS start distance. B, C) Quantification of soma-to-AIS start distance (B) and AIS length (C), under vehicle and KCl conditions, and KCl/vehicle ratio, at DIV21 (left) and DIV49 (right). B) DIV21, n=48-64 pictures/genotype (756-1204 AIS/genotype, 10 wells) and DIV49, n=15-29 pictures/genotype (343-565 AIS/genotype, 10 wells). C) DIV21 n=50 pictures/genotype (6888-8367 AIS/genotype, 10 wells), DIV49 n=47-90 pictures/genotype (1951-9161 AIS/genotype, 10 wells). Data represented as mean ± SEM. Statistical analysis= GLMM with Gamma distribution and log link. post hoc = Benjamini-Hochberg. * p_adj < 0.05 vs. control (CTRL, blue); # p_adj < 0.05 vs. 1 (dashed line).

Consistent with normal development, all genotypes showed AIS shortening and a distal-to-proximal shift over time (Fig. S11) (32). Compared to the control, KCNQ2-LOF iNeurons exhibited a significant distal AIS shift at both timepoints under naïve conditions, while AIS length remained unchanged in all lines except for reductions in KCNQ2-G290D at DIV21, and KCNQ2-A294V at DIV49 (Fig. 10).

Following 48 hours of KCl treatment at DIV21, the same significant AIS differences in the *KCNQ2*-LOF iNeurons compared to the control were observed as in the naïve conditions (Fig. 10B-C). Furthermore, all genotypes showed AIS plasticity post KCl treatment, indicated by a KCl:Vehicle ratio significantly different from one (Fig. 10B-C); AIS length decreased significantly across all genotypes (p<0.001) (Fig. 10C), and AIS position shifted further distally in the control, KCNQ2-A294V and KCNQ2-G290D (p<0.001, p<0.01, p<0.05, respectively) (Fig. 10B). At DIV49 post KCl treatment, the distal shift in AIS location was reduced for all *KCNQ2*-LOF lines compared to controls but only reached significance in the KCNQ2-R560W and *KCNQ2*-A294V lines (Fig. 10C). Contrary to control neurons, *KCNQ2*-G290D and *KCNQ2*-R560W showed a significant AIS length reduction (Fig. 10C).

In summary, our data show that under naïve conditions, *KCNQ2*-LOF variants cause a persistent distal shift of the AIS in both early and more mature neurons. While AIS plasticity at DIV21 appears intact, DEE neurons show altered plasticity at DIV49, either due to constrained AIS relocation from an already distally shifted AIS or enhanced AIS shortening in response to chronic depolarization.

## Discussion

In this study, we investigated the impact of KCNQ2-LOF variants on human cortical network development using glutamatergic iNeurons from individuals with KCNQ2-DEE (R560W, G290D, A294V) and KCNQ2-SeLFNE (K327Q). Our longitudinal multimodal approach revealed that the pathophysiology of KCNQ2-LOF in glutamatergic neurons is a dynamic process shaped by the primary loss of Kv7.2 function, compensatory adaptations and subsequent maladaptive changes.

To our knowledge, this is the first study to apply GAM to analyze longitudinal MEA data. By leveraging the flexibility of GAMs, we were able to model complex, non-linear changes in neuronal network activity over time, revealing a biphasic profile of network dysfunction. We first show that early in maturation, KCNQ2-LOF neurons display an accelerated onset of spontaneous firing and increased excitability. At the network level, KCNQ2-DEE lines also exhibited marked hyperexcitability, consistent with the findings of Simkin et al., who described the functional effect of the KCNQ2-DEE R581Q variant in iNeurons. We hereby validate that this is a variant-independent LOF effect (18).

Importantly, the most pronounced network-level differences in KCNQ2-DEE iNeurons emerged during the earliest stages of network burst formation. Our transcriptome data showed an increasing *KCNQ3*:*KCNQ2* expression ratio over time, aligning with previous findings that *KCNQ2* is expressed earlier and at higher levels than *KCNQ3* during brain development (14–17). If this transcriptional pattern translates to protein levels, it will favor the preferential formation of Kv7.2 homotetramers during early maturation, making this developmental stage particularly sensitive to *KCNQ2*-LOF. Interestingly, while the raster plots of Simkin et al. show minimal activity and no evidence of network bursting between DIV12–DIV15, our recordings reveal the emergence of network bursts as early as DIV14. This likely reflects the greater sensitivity and resolution of our HD-MEA technology, underscoring its utility in detecting important early neurodevelopmental phenotypes that may be missed with conventional low-density MEA.

We showed that these early network differences were Kv7-driven, as RTG application effectively reversed these effects in KCNQ2-G290D cultures. This is further supported by prior studies showing that acute pharmacological blockade of Kv7 channels in control iPSC-derived neuronal cultures induces a similar hyperexcitable network phenotype to that observed in our KCNQ2-LOF lines (33, 34). At this maturation stage, we observed that loss of Kv7 function altered expression of potassium channels and modulators, with an overall shift toward facilitating potassium efflux, likely as a compensatory mechanism to maintain potassium homeostasis. This adaptive mechanism may explain why total potassium current density did not differ between *KCNQ2*-G290D iNeurons and its isogenic control neurons. Additionally, all *KCNQ2*-LOF lines exhibited a distal shift in AIS positioning, mirroring effects seen in rodent neurons after chronic M-current suppression (27). However, these compensatory mechanisms, including relocation of the AIS and the rescue of the potassium homeostasis, appear not sufficient to overcome the *KCNQ2*-LOF induced hyperexcitability.

As maturation progresses, network parameters in *KCNQ2*-LOF cultures exhibited a marked shift, with several previously elevated metrics, such as burst duration, spikes per burst, burst ISI, falling below control levels. This suggests the involvement of time-dependent compensatory and potentially maladaptive mechanisms. Supporting this, RTG treatment during late maturation did not only fail to normalize these biphasic parameters but exacerbated them. The absence of a similar network-level transition in the study of Simkin et al. likely stems from their shorter MEA recording window (up to DIV31), which precedes the critical DIV30–50 window during which our *KCNQ2*-DEE cultures transitioned. However, Simkin et al. did report temporal changes at the single-cell level. Specifically, at DIV21, KCNQ2-DEE R581Q action potential characteristics could be partially reproduced by acute Kv7 channel blockade in control neurons, but by DIV35, only chronic Kv7 inhibition, and not acute, mimicked the disease phenotype (18). This suggests that compensatory changes emerge at the cellular level before manifesting as network level alterations.

The biphasic pattern was also evident at the synaptic level, reflected in both transcriptional and structural changes. During early maturation, synaptic GO enrichment was driven by upregulated genes linked to increased presynaptic glutamate release and enhanced postsynaptic receptor activation, consistent with the prolonged bursts and increased excitability observed in MEA recordings. In line with this, previous studies have shown that acute pharmacological blockade of M-channels increases neurotransmitter release (35, 36). In contrast, at the later maturation phase, synaptic enrichment was predominantly driven by downregulated genes indicative of reduced synaptic vesicle release. Disrupted synaptic vesicle recycling and reduced vesicle release have previously been linked to chronic epilepsy pathophysiology (37–43). Supporting this, Birtoli and Ulrich demonstrated that shorter IBI and thus increased burst frequency, can reduce the synaptic transmitter release due to insufficient recovery time between bursts, thereby shortening the duration of subsequent bursts, in line with the later-stage network characteristics we observed (44). Functional validation of these transcriptomic signatures in a follow up study would be valuable to confirm whether the observed gene expression changes effectively translate into corresponding alterations in synaptic function. This is particularly relevant as we also observed structural evidence of synaptic alterations: KCNQ2-LOF lines showed a significantly steeper decline in presynaptic density during maturation compared to controls.

Increased burst frequency is a recognized hallmark of network hyperexcitability that has been reported in several other iPSC-derived epilepsy models (18, 33, 45, 46). In our KCNQ2-LOF lines, elevated burst frequency may result from direct loss of Kv7, as this parameter responded favorably to RTG. However, additional postsynaptic mechanisms may also contribute. Specifically, transcriptomic data revealed reduced expression of the major SK channel, KCNN1, at DIV49, coinciding with reduced spikes per burst. In contrast, Simkin et al. reported a maladaptive upregulation of SK channels at DIV35, a stage when spikes per burst were still elevated, corresponding to the early phase of our model. This suggests that SK channel expression may follow a dynamic trajectory, shifting from early compensatory upregulation to later downregulation. Supporting this, Simkin et al. showed that pharmacological SK channel blockade with apamin reduced spikes per bursts, mirroring the late-stage phenotype we observed.

Although our study includes only a single SeLFNE-line (K327Q) without an isogenic control, limiting the strength of our conclusions, our findings suggest potential differences in network behavior and plasticity between SeLFNE and DEE-associated *KCNQ2* variants. During early maturation, when Kv7.2 homomers are likely the predominant Kv7 channel, the electrophysiological profile of the SeLFNE culture resembled that of the DEE-lines but showed only subtle, largely non-significant network alterations that transitioned earlier. Furthermore, when assessing activity-dependent AIS plasticity at DIV49 using chronic KCl treatment, impairments were observed exclusively in the DEE lines, while the SeLFNE line responded similarly to controls. Although these observations require cautious interpretation due to the limited sample size and absence of isogenic controls, they support the hypothesis that more pronounced early network disruptions, primarily driven by homomeric Kv7.2 homo-tetramers rather than Kv7.2/Kv7.3 hetero-tetramers, may compromise the network’s long-term adaptive capacity and may be critical for the severity of KCNQ2-DEE. This aligns with previous work demonstrating that the current density of homomeric channels in heterologous cell systems is the most reliable biophysical correlation of the DEE phenotype (47). In this regard, it is relevant to note that at later neurodevelopmental stages, spontaneous electrophysiological recordings no longer clearly differentiated DEE from SeLFNE cultures, suggesting that spontaneous MEA profiling alone may not fully capture the neurodevelopmental deficits inherent to DEE. One possibility is that these deficits in KCNQ2-DEE cultures may only become evident under specific conditions or functional challenges, as demonstrated by our activity-dependent AIS plasticity experiments. It is also important to consider that KCNQ2 is expressed not only in excitatory neurons but also in inhibitory neurons and astrocytes (48–50). For example, selective loss of KCNQ2 from GABAergic interneurons has been shown to increase network excitability in the immature mouse brain (51–53). Therefore, more complex *in vitro* models, such as co-culture systems incorporating inhibitory neurons and astrocytes, or organoid-based models may be necessary to distinguish DEE from SeLFNE phenotypes.

Finally, while acute RTG treatment at the late stage exacerbated several disturbed network parameters, this does not necessarily contradict the preliminary favorable clinical response to RTG seen in a small cohort of individuals with KCNQ2-DEE (10). Acute application may be insufficient to rescue long-term adaptive mechanisms, and chronic treatment of cultures is likely required to more accurately evaluate the therapeutic potential of RTG. This notion is supported by prior studies reporting differential effects of acute versus chronic RTG treatment on neuronal network behavior (54).

In summary, our longitudinal multimodal approach enabled us to map the dynamic impact of KCNQ2-LOF on cortical network development. We identified a biphasic pattern of dysfunction: an early, Kv7.2-driven hyperexcitability, followed by later maladaptive changes shaped by compensatory functional, transcriptional, and structural adaptations. Our findings highlight the importance of developmental timing in disease modeling and therapeutic testing, and underscores the value of human in vitro models for capturing the evolving nature of neurodevelopmental disease mechanisms.

## Acknowledgements

All authors listed have made a substantial, direct, and intellectual contribution to the work. The authors thank the patients, control individuals and their parents who volunteered after informed consent to be sampled to generate iPSC lines for research purposes. We thank Dr. Julia Faura Llorens and Matilde Malcorps for helpful discussions regarding analysis strategies for the RNA sequencing data. We thank Professor Nael Nadif Kasri and Jack Parent for the initial training of ND in neuronal culturing techniques and for their ongoing support and advice during troubleshooting.

## Author contributions

All authors listed have made a substantial, direct, and intellectual contribution to the work. ND, MK and SW designed the experiments. ND generated all NGN-iPSC lines. ND and MK performed all the MEA seedings and analysis. ND, MK, LC, NZ, EDV, VS, and EV performed the MEA recordings. ND performed the imaging and transcriptome experiments. BA and PV contributed to the imaging data acquisition and analysis. MK performed the pharmacological MEA experiment. GC, FM and MT designed and performed the patch clamp experiments. ND, MK, GC, FM and SW wrote the manuscript. All authors revised the manuscript and approved the final manuscript.

## Material and methods

### Patient inclusion and generation of human induced pluripotent stem cells

Experiments involving human-derived cells were performed after informed consent and approval by the Committee of Medical Ethics of Antwerp University Hospital and University of Antwerp (reference 19/20/257). Blood samples were taken from one proband with *KCNQ2*-SeLFNE (*KCNQ2*-K327Q), three individuals with *KCNQ2*-DEE (*KCNQ2*-G290D, *KCNQ2*-A294V, and *KCNQ*-R560W), and a healthy sibling of *KCNQ*-R560W (WT). The iPSC lines were generated by reprogramming subjects’ PBMCs by the RadboudUMC Stem Cell Technology Center (SCTC) using episomal vectors with the Yamanaka transcription factors *Oct4*, *c-Myc*, *Sox2* and *Klf4* (55). As additional control line a CRISPR Cas9 corrected clone of the *KCNQ2*-G290D iPSC line (ISO_G290D), generated by The Stem Cells Core Facility of the Institut du Cerveau et de la Moelle Epiniere – ICM in Paris, France, was added to the cohort of this study. The iPSC lines were tested for pluripotency markers (Nanog, Sox2 and Oct4) using qPCR, immunohistochemistry and trilineage differentiation, and showed no abnormalities on CNV analysis. Details for each iPSC lines are available in the Human pluripotent stem cell registry (https://hpscreg.eu/).

### iPSC maintenance and stable Neurogenin (NGN)-iPSC generation

The iPSCs were maintained in StemFlex medium (Thermo Fisher Scientific, A3349401) on Geltrex (Thermo fisher Scientific, A1413302) coated plates at 37°C and 5% CO2. Cells were passaged using ReleSR (Stemcell Technologies, 05872) when they reached 70% confluency. iPSC lines were checked for mycoplasma contamination every three months.

Induced overexpression of human Neurogenin (hNGN) 1 and 2 was used to generate iPSC-derived cortical excitatory neurons (see workflow in Fig. 1A). Stable NGN-iPSC lines were generated using two TALEN plasmids (pZT-C13-R1 and pZT-C13-L1 Addgene ID: #52638 and 52637, respectively) targeting the CLYBL locus to integrate a construct containing hNGN1 and hNGN2 under a doxycycline inducible promoter, as well as an EF-1α promoter driving constitutive expression of mCherry (pUCM-CLYBL-Ngn1/2, a gift of Dr. Michael Ward), as described by Tidball et al. 2020. In brief, iPSCs were dissociated using Accutase (Sigma, A6964) and a 1E6 cell pellet was resuspended in 20 µl supplemented Nucleofector solution of the P3 Primary Cell 4D-nucleofector® X Kit S (Lonza, V4XP-3032) and 2 µl of plasmid mix (1:1:1, 1 µg/µl). The suspension was nucleofected with the 4D-Nucleofector Core and X Unit (Lonza), using program CB-150. Ten minutes post-nucleofection, cells were plated in StemFlex media supplemented with 10 µM Y-27632 rho kinase inhibitor (Tocris, 1254). iPSC lines were expanded to a full 6-well plate. They were dissociated and sorted using the mCherry signal to enrich for successfully engineered iPSCs. The dissociated iPSC pellet was resuspended in 1 ml FACS buffer (PBS with 2% FBS), with 1000X Viability Dye eFluor® 780 (Thermo Fisher Sci., 65-0865) and incubated for 20 min on ice. After incubation, 10 ml FACS buffer was added, followed by centrifugation for 3 min at 300x g to wash the cells. The cell pellet was resuspended in 2 ml FACS buffer and ran through a 30 µm filter. The cell suspension was added to the MACSQuant® sorting cartridge (Miltenyi Biotec). Gating hierarchies were constructed using MACSQuant® Tyto software before sorting. Cell debris, doublets, and dead cells were gated out. The mCherry+ sorted cells were collected from the cartridge and plated at low density to generate clonal lines. 250 and 500 cells were separately seeded onto Geltrex-coated 6 well plates in

StemFlex media supplemented with CloneR™ (stem cell technologies, 05888). Three hours post-seeding, cells were placed in an Incuctye® S3 Live Cell Analysis System and imaged every 3 hours for 7 to 10 days to track colony formation from single cells. Cells were kept in CloneR supplemented media for 4 days, after which the CloneR supplement was removed. Between day 7 and day 10, colonies originating from single cells were picked and placed into individual wells of a 24-well plate coated with Geltrex and containing StemFlex media. Clonal lines were then split into duplicate wells of a 12-well plate; one for expansion and one for DNA isolation to check the zygosity of the NGN-construct. Zygosity validation was performed using a multiplex of 2 primer pairs (Table 6), one targeting the CLYBL locus around the insertion site, and the other within the integrated NGN-construct. All homozygous clones were checked for the recurrent duplication of chr20q11.21 (15, 16) using Multiplex Amplicon Quantification (MAQ). MAQ negative clones were subsequently analyzed for other CNVs using a single-nucleotide polymorphism (SNP) microarray analysis.

### Multiplex Amplicon Quantification

To exclude the presence of the recurrent duplication of chr20q11.21 (15, 16) in the corrected clonal lines, we used our in-house Multiplex Amplicon Quantification (MAQ) method (Agilent). This method includes multiplex PCR amplification of fluorescently labelled target and control amplicons, followed by fragment analysis. The assay targets six amplicons situated in and around the chr20q11.21 region, along with five control amplicons situated at random genomic positions outside the chr20q11.21 region and other known CNVs. These 11 amplicons were PCR-amplified in a single reaction using 20 ng of genomic DNA template. The peak areas of the target amplicons were normalized to those of the control amplicons. Comparisons of the normalized peak areas between clonal lines and reference samples were used to produce a dosage quotient (DQ) for each targeted amplicon, computed by the MAQ-S software package (Agilent). DQ values exceeding 1.25 were considered indicative of a duplication.

### Generation of iPSC-derived NGN neurons

NGN-iPSCs were dissociated using Accutase and 1E6 cells were seeded per well in a 6 well plate with 2 ml StemFlex supplemented with 10 µM Y-27632 rho kinase inhibitor (Tocris, 1254) and 2 µg/ml doxycycline (Sigma, D9891). After 24 hours, 1 ml of spent media was removed and 2 ml of StemFlex supplemented with 2 µg/ml doxycycline added. After 48 hours, 2 ml spent media was again removed and replaced with 2 ml of StemFlex supplemented with 2 µg/ml doxycycline. 72 hours after initial induction, these pre-differentiated day 3 neurons (D3N) became post-mitotic and were frozen in freezing media (90% KnockOut Serum Replacement (KOS), 10% DMSO). For final differentiation, D3N were co-cultured with primary mouse astrocytes in Complete Seeding media containing Neurobasal Plus basal media, B27 plus (50x), BDNF (10 ng/ml), NT3 (10 ng/ml), doxycycline (2 µg/ml) (Sigma, D9891), cytosine b-D-arabinofuranoside (Ara-C) and CEPT Cocktail (HY-K1043, MCE). From day 7 onwards, half-media changes were performed twice a week using complete differentiation media containing Neurobasal Plus basal media, B27 plus (50x), FBS (50x), BDNF (10 ng/ml) and NT3 (10 ng/ml).

### qPCR

Total RNA was isolated using the Macherey-Nagel NucleoSpin RNA mini kit (Macherey-Nagel, 740955) according to manufacturer’s protocol. cDNA was reverse-transcribed from total RNA using the iScript cDNA Synthesis Kit (Bio-Rad Laboratories, 1708890). qPCR reactions were performed in triplicate using SYBR Green Real-Time PCR master mixes (Applied Biosystems, 4309155). Expression levels were normalized to GAPDH and TBP, and the ΔΔCt method was used to determine the relative levels of mRNA expression (56). Primers for qPCR reactions are included in table 6.

### Dissection of mouse brains and generation of primary mouse astrocyte culture

All animal experiments were approved (ECD 2020-076) and conducted in accordance with the policies and guidelines set forth by the Ethical Committee for Animal Testing at the University of Antwerp. These policies and practices comply with European Committee Guidelines (Directive 2010/63/EU). Primary mouse astrocytes were isolated from postnatal day 0-3 C57BL/6, 129/Sv and 129/Sv-KCNQ2^274/+^ KI mice pup brains. In brief, the cortex was isolated from the mouse brain and the meninges removed. Cortices were cut into smaller pieces and dissociated in 0.25% trypsin at 37°C, 5% CO2 for 10 min. Trypsin was removed and the cell suspension was resuspended 10 times in glia media (DMEM, 10% FBS, 100x Glutamax, 50 U/ml P/S, 1 mM Sodium Pyruvate) supplemented with DNase I. The cell suspension was incubated statically for 5 min, whereafter the supernatant, containing the single cells, was transported to another conical tube. This was repeated 3 times. The cell suspension was centrifugated at 300x g for 3 min to remove the DNase I. The pellet was resuspended in glia media, filtered through a 0.45-micron filter, and seeded on Geltrex coated T75 flasks. When confluent after approximately 5-7 days, the flask was shaken vigorously for 30 seconds to remove all non-astrocyte contaminant cells Mouse astrocytes of passage 2 were checked for mycoplasma and frozen in 90% FBS/10% DMSO. One week before the neuronal seedings, frozen mouse astrocytes were thawed and plated in glia media on a Geltrex coated T75 to allow the astrocytes to recover from the freezing process.

### Patch clamp recordings

Whole-cell patch-clamp recording in voltage- and current-clamp configurations were performed at room temperature (22-24°C) using a commercially available amplifier (Axopatch 200B, Molecular Devices). Data were sampled at a frequency of 15 kHz, filtered at 5 kHz with the 4-pole low-pass Bessel filter of the amplifier, and digitized using a Digidata 1440A (Molecular Devices, USA). The pCLAMP software (version 10.0.2) was used for data acquisition and analysis. Recording pipettes (5-6 MΩ resistance) were pulled from borosilicate glass capillaries. Solutions were applied using fast-solution exchange perfusion system (CFPKGHB8, Green Leaf Scientific, Ireland). Series resistance was typically ≤ 15 MΩ, and, in most recordings, compensated by at least 80%. Neurons plated on glass coverslips were continuously perfused with artificial cerebrospinal fluid (aCSF) bubbled with a mixture of CO2 (5%) and O2 (95%). aCSF contained (in mM): 124 NaCl, 2.5 KCl, 26 NaHCO3, 1.25 NaH2PO4, 2 MgSO4, 2.4 CaCl2, 15 glucose, pH 7.35, osmolarity 305-310 mOsm. Recording pipettes were filled with standard intracellular solution containing (in mM): 125 K-gluconate, 25 KCl, 10 HEPES, 10 Na2-phosphocreatine, 4 Mg-ATP, 0.4 Na 345 GTP, pH 7.35 adjusted with KOH; osmolality 280-290 mOsm. No correction was made for liquid junction potential. Neurons meeting the following criteria were included in the analysis: holding current at−80 mV <100 pA, resting membrane potential (RMP) <-35 mV, series resistance < 20 MΩ, and input resistance > 250 MΩ. Resting membrane potential was determined by averaging for 20 s of recording. Afterward a small holding current (less than 5 pA) was used to clamp the resting membrane potential close to −65 mV. Current ramps were elicited from an initial current step of −50 pA for 0.5 s followed by depolarizing ramp from −50 to +200pA for 2 s. This protocol was used to calculate the action potential (AP) properties as described in Simkin et al. (18)

### HD-MEA plate preparation and seeding

Prior to cell plating, HD-MEA plates were pre-treated with 1 ml of 1% tergazyme for 1 hour, washed 3 times with PBS and sterilized in 70% ethanol for 30 min. After removing ethanol and washing, 1 ml of Neurobasal Plus basal media with B-27 Plus supplement (50x) was added per well and incubated for 48 hours (37°C) to condition the chips. Afterwards, the media was removed, and the chips were washed once with 1 ml PBS. The chips were coated first with 50 μl 0.14% PEI in borate buffer for 1 hour, then extensively washed four times with 1 ml Milli-Q water followed by airdrying overnight. The following day, the chips were coated with 20l1μg/ml CellAdhere™ Laminin-521 (Stemcell Technologies, 200-0117) in DMEM for 0.5 to 1 hour at 37°C. During this incubation time cell suspensions for plating were prepared. Frozen D3N were thawed at 37°C in a warm water bath, and in parallel, fresh mouse astrocytes of passage 2 or 3 were dissociated using 0.05% Trypsin. A mixture of 5E5 D3N and 5E4 astrocytes was added directly to the chips in a 50 μl cell suspension composed of Complete Seeding media supplemented with Primocin (100 μg/ml) and incubated for 30 min in a 5% CO2 cell culture incubator at 37 °C. After cell adherence, an additional 2 ml Complete Seeding media supplemented with Primocin was gently added in the wells.

### HD-MEA recordings

#### Activity and Network Assay

All MEA recordings were performed on a MaxTwo (MaxWell Biosystems AG, Zurich, Switzerland), using MaxLab Live assays as described previously by Ronchi et al. 2020. Activity Scan Assays and Network Assays were performed three times a week during critical maturation period (Monday, Wednesday, Friday) from DIV12 to DIV28, and twice a week (Monday, Friday) DIV28 to DIV49 at 5% CO2 and 37°C. For all the recordings, a 300 Hz high-pass filter was used and a spike detection threshold set to 5.0 standard deviations. For the Activity Scan Assay, the checkerboard configuration was used with a recording time of 30 sec for each configuration. For the Network Assay, 1024 electrodes were selected using the Maxlive neuronal selection algorithm which selects the electrodes with the highest amplitude while also maximizing the distance between the individual electrodes. This reduces the potential for redundant recording of the same cells with multiple electrodes and therefore increases the total number of measured cells. Network measurements were recorded for 10 min. All HD-MEA media changes took place immediately following recordings to allow >48 hours of media conditioning.

To account for the presence of long-tailed bursts until DIV28, we performed burst parameters analysis using two different peak-to-baseline cut-offs, influencing the start and stop detection of the burst, for data generated between DIV14 and DIV28 (Fig. 3). An 80% cut-off was used to analyze the period of high-frequency spiking that defined the main body of the burst, excluding the prolonged tail which results from residual network activity rather than primary bursting dynamics. A 95% cut off was used to include the tail in the burst, which is particularly important for outside burst parameters including inter spike interval (ISI) outside of the bursts and inter-burst interval (IBI), but also percentage spikes per burst, spikes per total burst (SPB total) and total burst duration (BD total) (Table 3). For data recorded after DIV28, only the 80% cut off was applied, as long-tailed bursts were no longer present, making tail inclusion unnecessary.

#### Pharmacological studies

Acute retigabine exposure was performed either on DIV21 or between DIV53 and DIV63. Baseline untreated recordings were obtained using the standard Activity and Network scan parameters as described above. Concentration-response measurements were performed by adding 10 μl of retigabine (10 mM stock in DMSO) or vehicle (% DMSO corresponding to equivalent exposure from associated retigabine concentration), in Complete Recording media, followed by 4 min diffusion, Network scan, and Activity scan, then progressing to the next ascending concentration. After reaching the maximal concentration, 20 µM of the Kv7.2 channel blocker XE-991 was added to each well, followed by recording as previously described. IC50 values were calculated using active area metrics normalized to untreated response. Semi-log normalized data were fit to IC50 curves using GraphPad Prism.

#### Immunostaining

Neurons cultured on 96-well glass bottom plates (P96-1.5H-N, CellVis) were fixed at DIV49 using 4% paraformaldehyde for 15 min. After washing with PBS, the cells were permeabilized with blocking buffer (PBS, 0.1% Bovine serum albumin (Sigma B4287), 10% Goat serum, 0.1% Tween-20) supplemented with 0.5% Triton X-100 for 30 min. Cultures were incubated with the primary antibody overnight at 4°C in blocking buffer. After two PBS washes, the cells were incubated with the secondary antibody in blocking buffer for 2 hours at room temperature. Following another PBS wash, Hoechst (5000x) was added for 10min. After a final wash, plates were topped with 300ul PBS and sealed. Images were acquired using a high-content microscope, a Nikon spinning disk confocal (Ti2 with W1 disk), using a 40X NA0.95 objective and standard filters for DAPI, FITC, Cy3 and Cy5. Antibodies used in this study are listed in Table 7.

#### AIS imaging analysis

To assess neuronal plasticity of the AIS, neuronal cultures were treated with either 10mM KCl or Vehicle for 48 hours. Two independent differentiation rounds with five wells per round for each condition were stained for both DIV21 and DIV49. For a comprehensive representation of the neuronal culture, ten images were taken per well. To measure the distance of the AIS to the soma, straight lines were manually drawn from the start of the AIS (marked by anti-Ankyrin-G staining) to the start of the soma (marked by anti-MAP2 staining) for each neuron using the line tool in ImageJ. The AIS length was measured exclusively from the anti-ankyrin-G channels. Images were processed using Ilastik (version 1.4.0) pixel- and object-level classification to identify the AIS. Output files of ImageJ and Ilastik were further processed using R for statistical analysis and graphical visualization.

#### Synaptic density analysis

iNeuronal cultures were stained at DIV21 and DIV49 using, pan-presynaptic marker, Bassoon, glutamatergic-specific post-synaptic marker, Homer1, neurite marker, MAP2 and DNA marker, Hoechst. Fifteen random Z-stacks (1 µm spacing) were acquired in each well, which originated from 2 independent differentiation rounds (10 wells per round). Z-projected images, for synaptic spot quantification, were analyzed in FIJI, and data in R, using in house developed scripts (available from https://github.com/DeVosLab/NeuroConnectivity) (57, 58). Briefly, nuclei were detected in the DAPI channel using the Stardist algorithm (59), and the neurite mask was construed using the MAP2 channel. Within a search area, i.e., the dilated neurite mask minus the nuclei mask, both pre-(Bassoon) and postsynaptic (Homer) synaptic spots were detected using Laplacian filtering. The criterion for overlap was set at least 5 overlapping pixels across both binarized images. Resulting numerical data was plotted in R, with one data point representing 1 well.

#### RNA sequencing

iNeuronal cultures of four replicates per cell line were harvested at DIV21 and DIV49. RNA fraction was collected using Qiazol and purified using the RNA MN kit according to the manufacturer’s instructions. The quality of the RNA was assessed with Fragment Analyzer 5200 (Agilent, Santa Clara, CA), using the DNF-471 RNA Kit. Samples were loaded according to manufacturer instructions. RNA quality numbers (RQN) values were calculated using the PROSize software (version 3.0.1.6). RQN values ranged between 9.7 and 10. Strand-specific transcriptome resequencing was performed by BGI (Shenzhen, China) using the DNBSEQ platform with PE100. Analysis was performed in-house with a standardized pipeline integrated in Genomecomb (60). The pipeline used fastq-mcf (http://code.google.com/p/ea-utils) for adapter clipping. Reads were aligned to the hg38 genome reference using STAR (61) in 2-pass mode and the resulting SAM file converted to CRAM format using Samtools (62). BAM files were sorted. Variants were called at all positions with a totalcoverage >= 5 using both GATK HaplotypeCaller (63) and Strelka (64). At the initial stage positions with a coverage < 8 or a genotype quality score < 25 were considered unsequenced. Gene counts were obtained using RNA-SeQC (65) with GENCODE(66) as the reference gene set.

#### RNA sequencing analysis

Differential gene expression analysis was performed using the DESeq2 package (version 1.40.2) in R. Genes with fewer than 8 counts across all samples were excluded from downstream analysis. For differential gene expression, count matrices and corresponding sample metadata were used to construct DESeqDataSet objects in DESeq2. Three distinct experimental designs were implemented to address the specific structure of each comparison using DESeqDataSetFromMatrix function. For the full dataset, a multi-factor design was specified to assess the effects of genotype, time, and their interaction. For individual time points, separate datasets were constructed for DIV21 and DIV49, assessing the effects of genotype. Normalization and variance stabilization were performed as implemented in DESeq2. Principal component analysis was performed on scaled data, with the prcomp function. DEGs were identified using Benjamini-Hochberg adjusted p-value < 0.05 and absolute log2 fold-change ≥ 0.58 (corresponding to a fold change of 1.5). To identify biological processes significantly enriched among significant genes, we performed Gene Ontology (GO) enrichment analysis using the clusterProfiler package. Enrichment was adjusted using the Benjamini-Hochberg method (FDR < 0.05). Synapse-specific gene set enrichment was conducted separately on downregulated and upregulated DEGs at DIV21 and DIV49, using the SynGO tool (https://www.syngoportal.org). To evaluate the enrichment of epilepsy-related and cardiac disease-related genes, Fisher’s Exact Test was performed. The epilepsy gene set was obtained from the Genes4epilepsy database (September 2024 release(67)), and the cardiac disease-related gene set was sourced from the T2Diacod database (t2diacod.igib.res.in/cardio). These gene sets were compared to significantly differentially expressed genes identified from transcriptomic analyses. For each comparison, a 2×2 contingency table was constructed to represent the overlap and exclusive members of the gene sets and the differentially expressed genes. Enrichment was assessed using Fisher’s Exact Test.

#### Statistical analyses

All analyses were performed in R version 4.3.0, or GraphPad Prism 10.4. In all figures, *P*-values are indicated as follows: \**P* < 0.05, \*\**P* < 0.01, \*\*\**P* < 0.001. In each figure legend, the n = (a)/(b) indicates the number of wells, pictures or cells (*a*)/ differentiation rounds (*b*). All results containing only one or two timepoints were analyzed using either linear mixed-effects models (LMMs) or generalized linear mixed-effects models (GLMMs), depending on the distribution of the response variable. Normality of residuals was tested via Shapiro–Wilk tests; LMMs (using the lmer function from the lme4 package) were applied to normally distributed data, while GLMMs (using glmer) with a Gamma distribution and log-link were used for positively skewed data. In both model types, fixed effects included experimental group, condition (e.g., maturation stage or compound), and their interaction, while differentiation round was included as a random effect to account for inter-experimental variability. Estimated marginal means and pairwise contrasts were computed using the emmeans package (Lenth, 2023). Both within-group and between-group contrasts were assessed depending on the experimental design. P-values were obtained for all relevant comparisons and adjusted for multiple testing using the Benjamini–Hochberg procedure.

To analyze the non-linear trends of the MEA measurement over time, we employed generalized additive modeling (GAM). GAM is a flexible statistical technique that, like generalized linear models, can handle incomplete observations and incorporate fixed and random effects. This approach extends traditional models by allowing the fitting of non-linear relationships between predictors and the response variable, making it particularly effective for accurately modeling non-linear longitudinal data. GAMs were fit to each parameter of interest using the gamm4 and mgcv package in R. For each parameter, the following GAM structure was used:

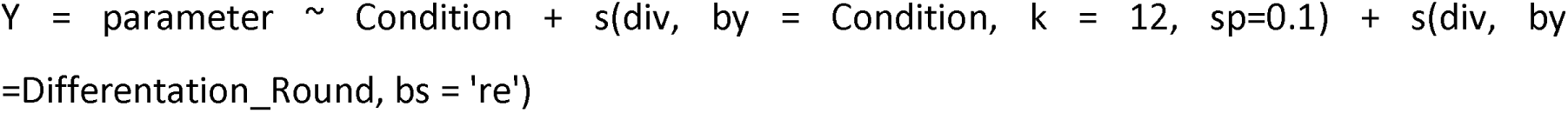

Condition indicates the experimental group, and the smooth term s(div) is used to capture non-linear changes over time. The by term enables modelling of time-specific trends for each Condition, and random smooth (e.g., bs = ‘re’) was applied to account for random effects of differentiation round. A fixed smoothing parameter (sp = 0.1) was applied to control the degree of smoothness in the time trend.

After generating the GAM model for each mutant line, pairwise comparisons between the smooth functions of mutant and control lines were performed, along with the corresponding simultaneous confidence intervals (SI) as described by Mundo et al. Regions where the SI did not cover zero were considered significantly different. To improve visualization of all the parameters of the Activity Scan Assay or the Network Scan Assay over time, heatmaps of the significant differences (SD) and the fold changes (FC) were generated using the GAM output.

### Data availability

The generated data are available from the corresponding author upon reasonable request.

