## Supplementary Figures 1-11 for "KCNQ2 Loss-of-Function variants disrupt neuronal maturation via early hyperexcitability followed by maladaptive network remodeling"

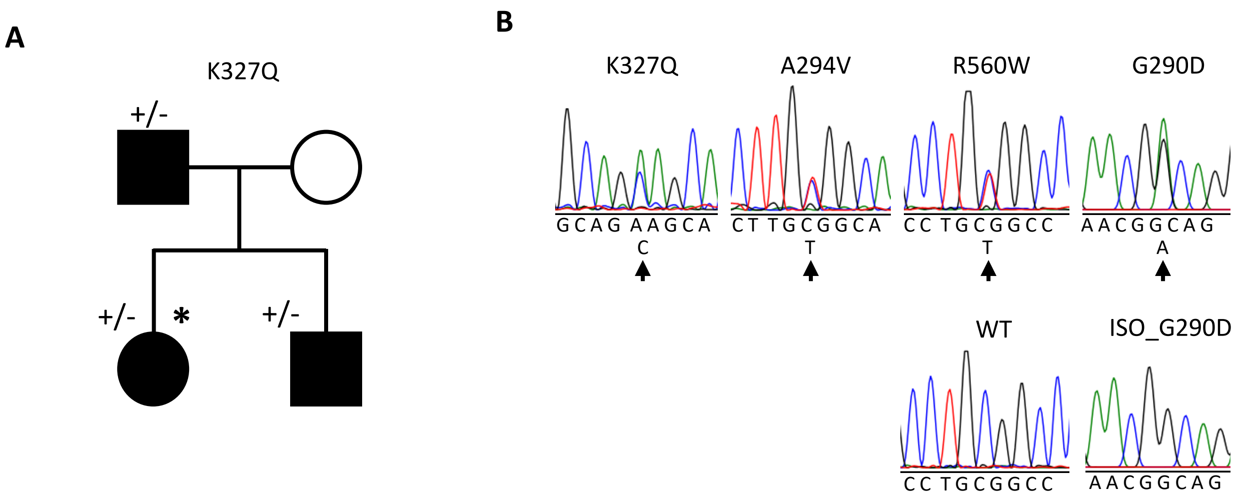


**Fig. S1 Pedigree and Sanger sequencing traces of KCNQ2 variants**. Left) Pedigree of the SeLFNE family carrying the segregating KCNQ2-K327Q variant (+/-). Astrix indicates individual from whom iPSC line has been derived. Right) Sanger traces of all KCNQ2 LOF variants (black arrow) and isogenic correction of G290D with PAM mutation (red arrow).

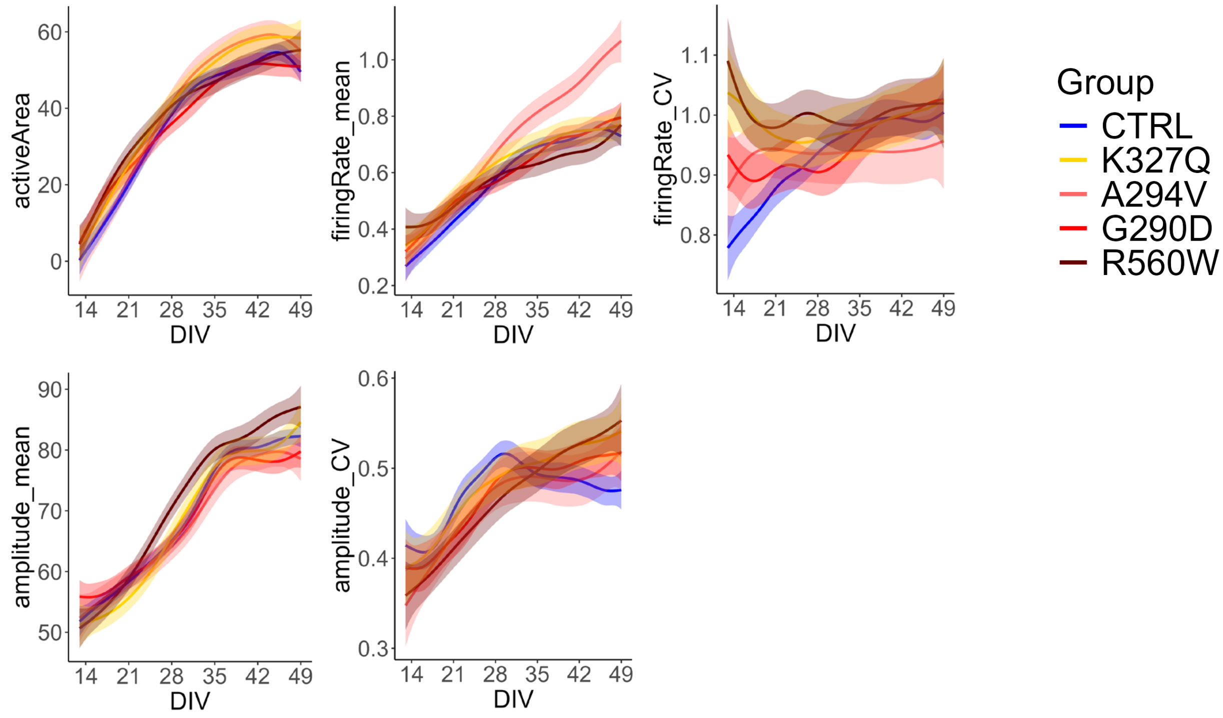


**Fig. S2 Smooth estimates of activity scan parameters of HD-MEA across genotypes over time.** Generalized additive models (GAMs) were used to estimate the smooth for all genotypes over days in vitro (DIV). Each panel represents a different electrophysiological parameter. Solid lines indicate the estimated mean trajectory, while shaded areas represent the 95% confidence intervals.

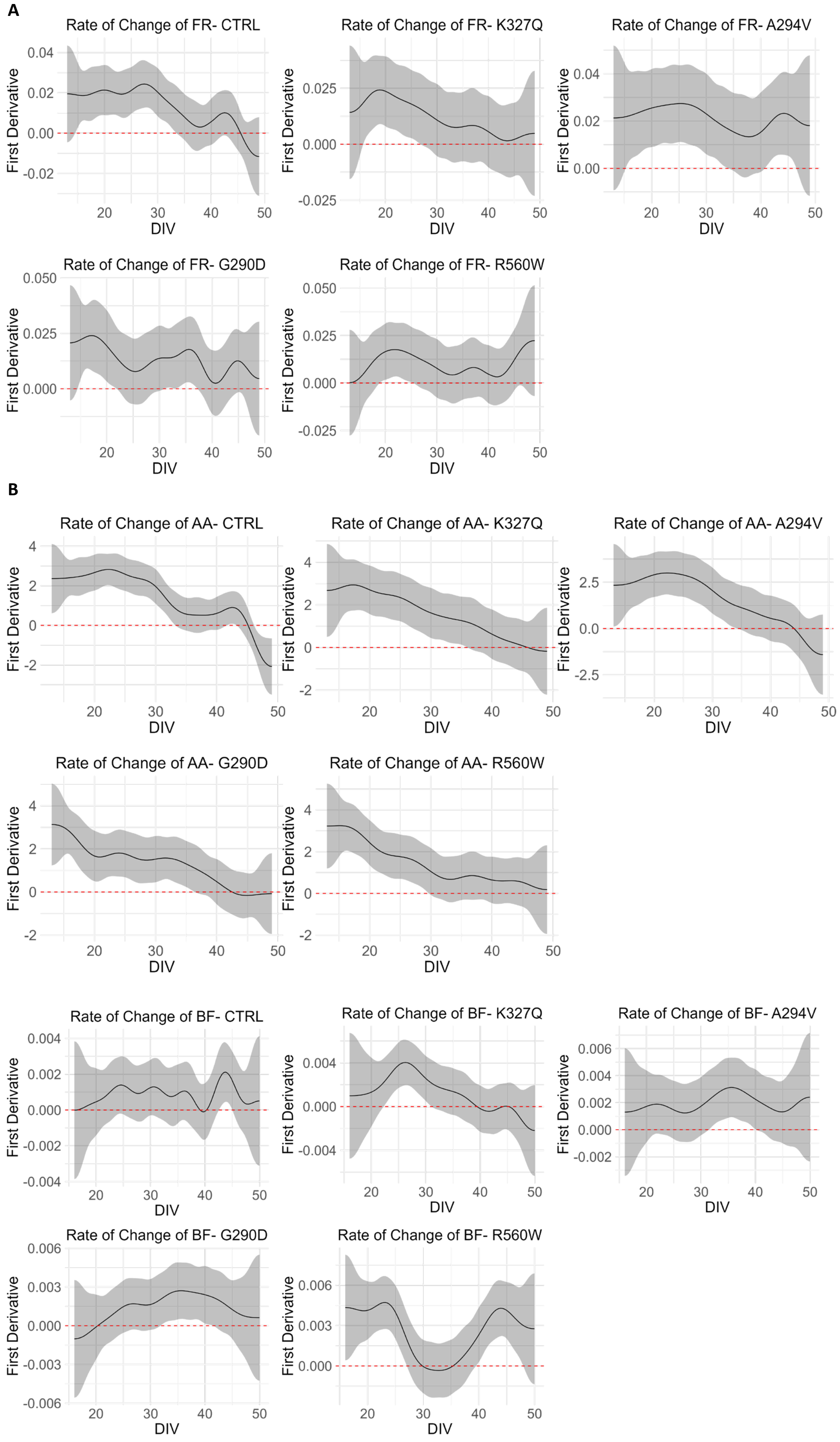


**Fig. S3 Directional Changes in HD-MEA derived Activity and Network Parameters Over Time**. The first derivative of (A) firing rate (FR) and (B) active area (AA) and (C) burst frequency (BF) over days in vitro (DIV) is shown for each genotype. Solid lines indicate the estimated first derivative, while shaded areas represent the 95% confidence interval. The red dashed line at y=0 denotes the threshold between increasing and decreasing trends. Positive values indicate a net increase over time, while negative values indicate a decline

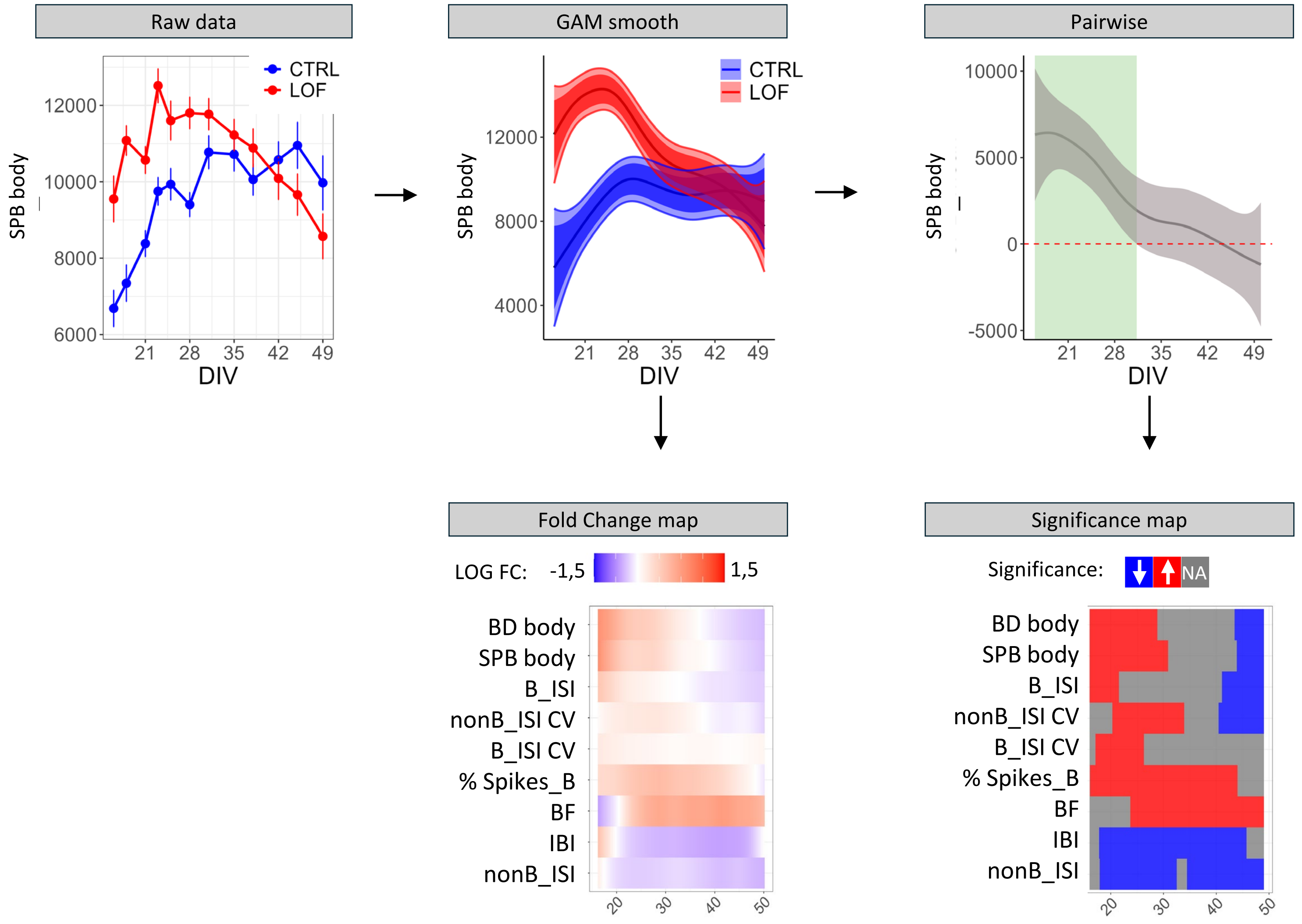


**Fig. S4 Visual presentation of the GAM pipeline**. RAW data of all MEA parameters are first modelled as GAM smooths. From the smooths the fold change and pairwise difference between mutant and control is extracted. Regions where the simultaneous confidence intervals did not cover zero were considered significantly different (green region in pairwise). All significant regions are visualized in a significance heatmap. Positive pairwise values and fold changes indicate an increase in the mutant line for the respective parameter compared to wildtype and are represented in red, while negative values suggest a decrease and are presented in blue.

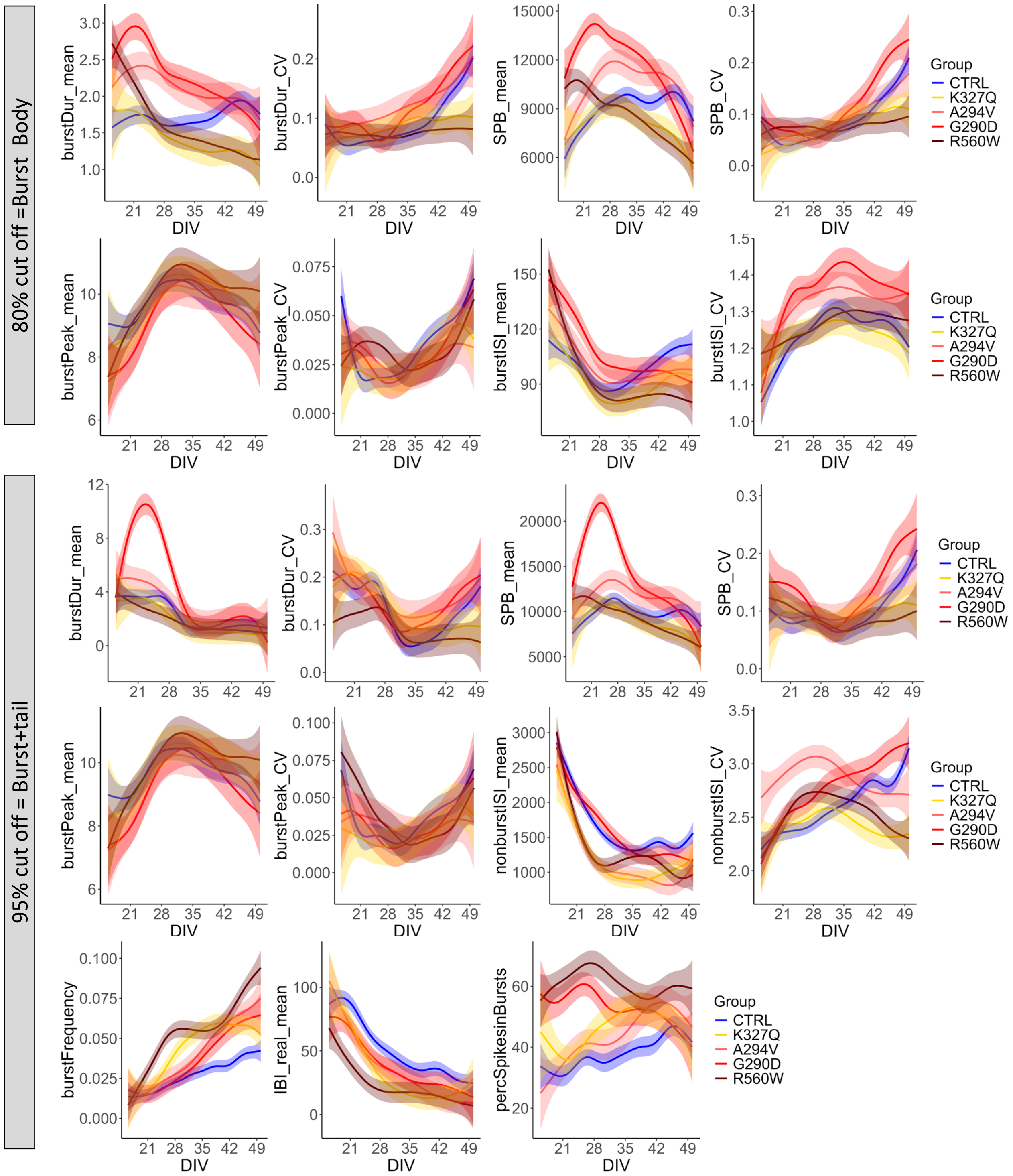


**Fig. S5 Smooth graphs representing the development of network scan parameters over time.** Generalized additive models (GAMs) were used to estimate the smooth trajectories for all genotypis over days in vitro (DIV). Each panel represents a different electrophysiological parameter. Solid lines indicate the estimated mean trajectory, while shaded areas represent the 95% confidence intervals.

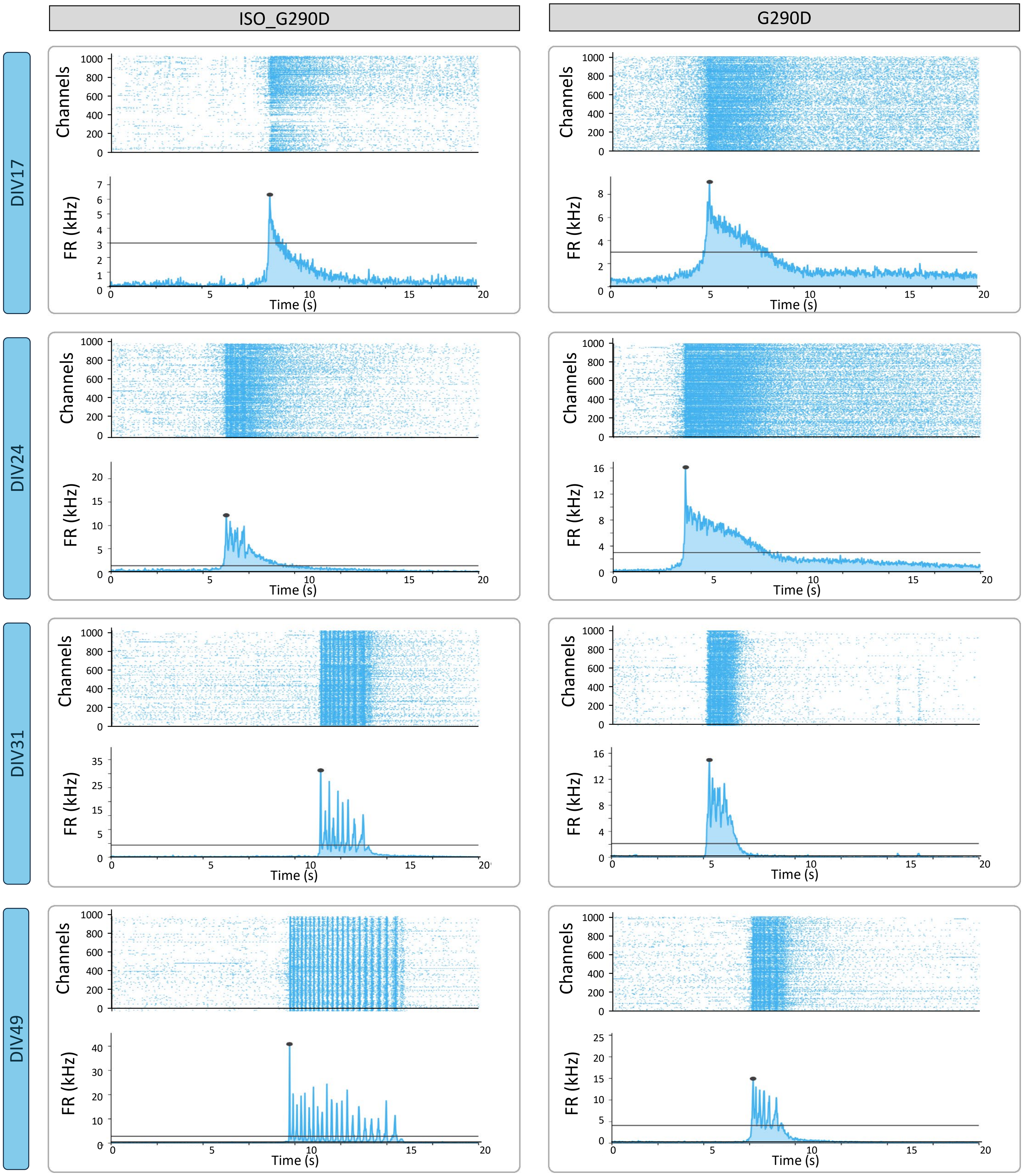


**Fig. S6 Visual representation of the dynamic changes in burst wave form during development and between genotypes**. Representative burst wave form images of iNeurons of KCNQ2-DEE G290D and ISO_G290D at four different time points; DIV17, DIV24, DIV31, DIV49. Raster plot and network plot are shown for each timepoint,

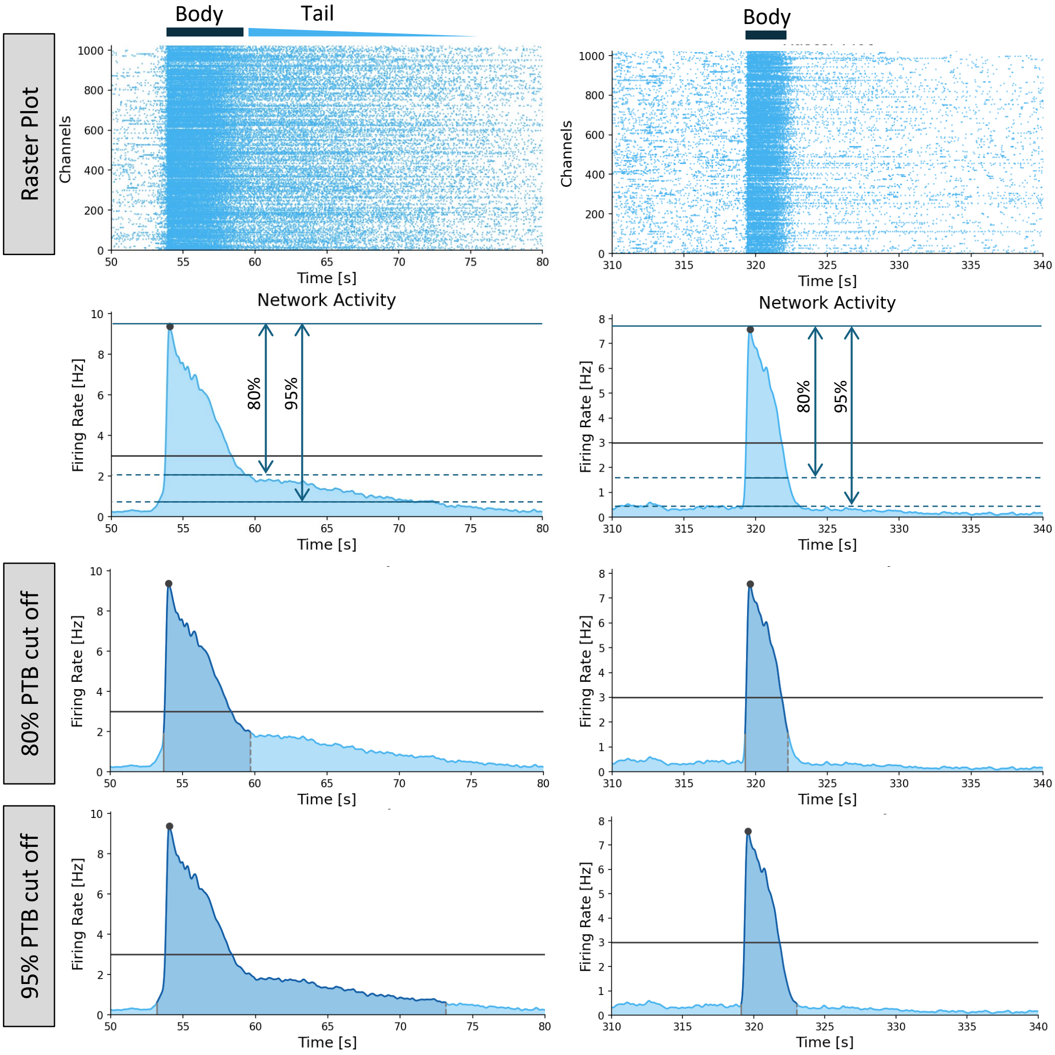


**Fig. S7 Overview of two-way analysis for burst parameters in early development**. Top) Raster plot of (left) KCNQ2-G290D and (right) WT iNeurons at DIV21, showing the presence and absence of a burst tail respectively. Bottom) Burst wave form visualization using firing rate with a smoothing window size of 0.1 seconds. An 80% and 95% Peak-to-Baseline cut off was used to extract burst parameters that capture the burst body and burst body plus burst tail respectively. Solid black lines indicate the firing rate threshold for burst detection. Detected burst are indicated with a black dot at the peak.

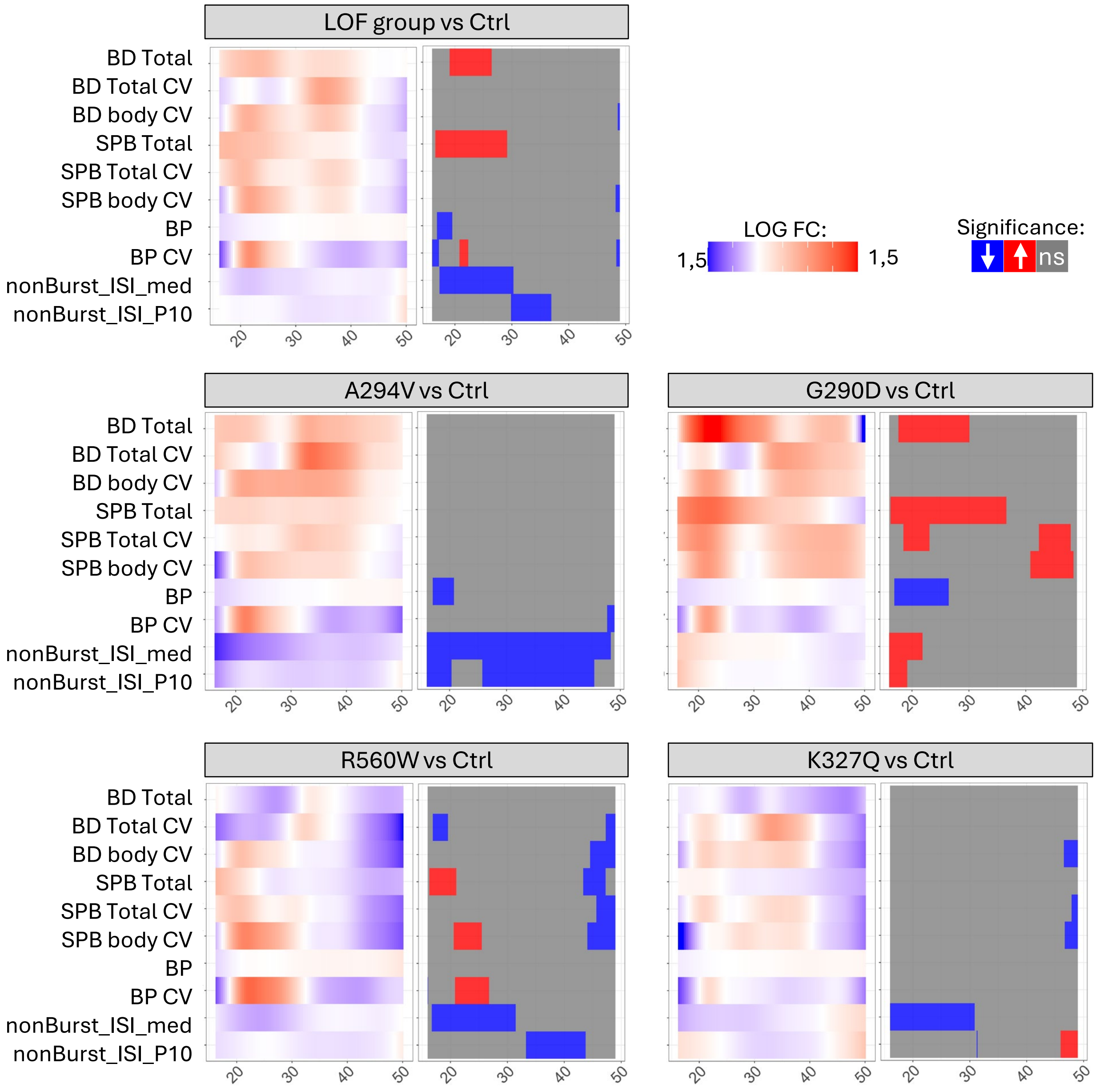


**Fig. S8 Longitudinal comparison of non-shared HD-MEA network parameters of KCNQ2-LOF iNeurons to its control.** A) Grouped LOF data; B) Genotype-specific data. Comparative Heatmaps show fold-change (FC) vs. CTRL; pairwise comparisons indicate significance. Positive FC and pairwise values (red) = increase; negative (blue) = decrease. n=39/11 CTRL, 16/6 K327Q, 13/6 A294V, 19/5 G290D, 19/7 R560W. BD: Burst duration, SPB: spikes per burst, BP: Burst Peak, nonB_ISI_med: median burst inter spike interval outside of burst, CV: coefficient of variance, P10: 10 percentile.

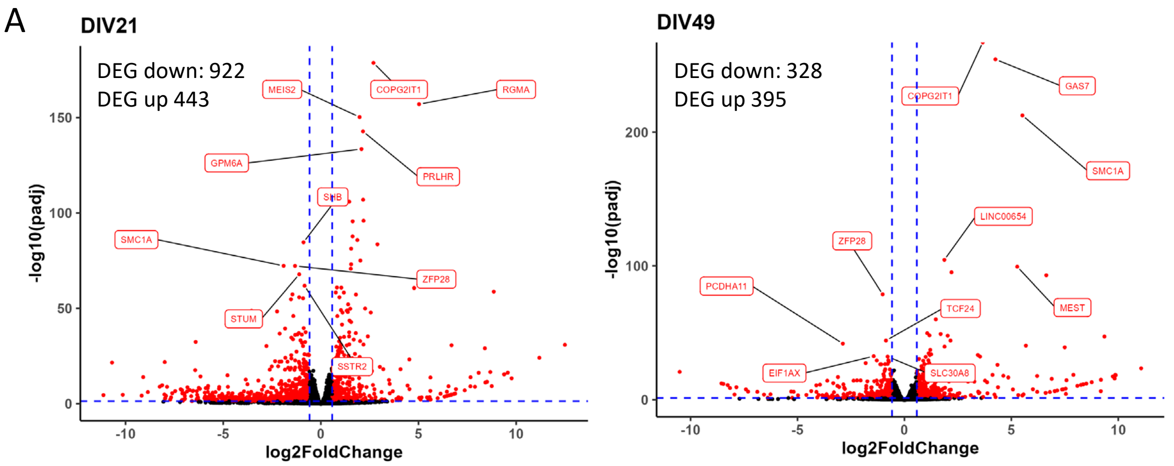


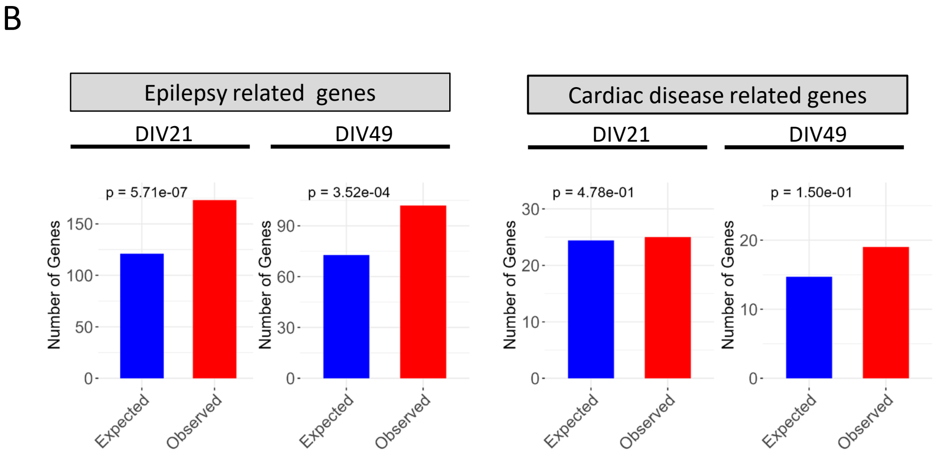


**Fig. S9 Differential expression and enrichment analysis in KCNQ2-G290D iNeurons.** A) Volcano plots of DEGs (FDR < 0.05, |logFC| > 0.58) at (A) DIV21 (1,365 DEGs) and (B) DIV49 (723 DEGs). Top 5 most significant genes are labelled. Dashed lines indicate significance thresholds. B) Enrichment analysis comparing observed vs. expected overlap of significant genes with (left) epilepsy-associated genes and (right) cardiac disease-related genes. Bars represent the observed overlap (blue) and expected overlap by chance (red). P values are calculated using Fisher's exact test.

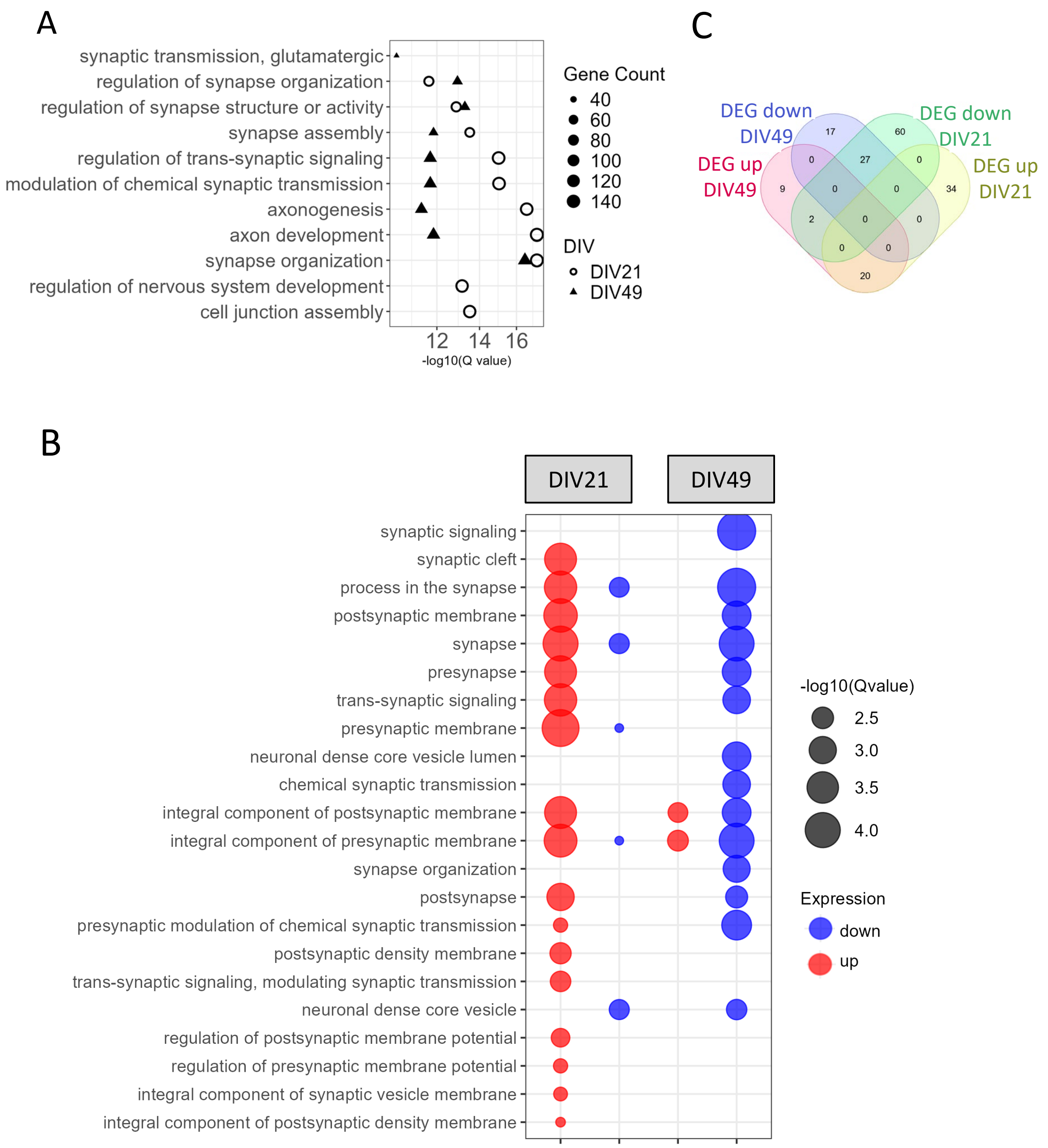


**Fig. S10 General and synaptic specific *Gene Ontology (GO)* enrichment analysis in KCNQ2-G290D iNeurons.** A) *Top 10 enriched GO biological process terms for significantly expressed genes (FDR < 0.05) at DIV21 and DIV49, based on individual analyses per timepoint. (B) Venn plot of overlapping synaptic DEGs between DIV21 and DIV49, with directionality of expression indicated. C) Synaptic GO term enrichment (SynGO) performed separately for upregulated and downregulated differentially expressed genes (DEGs) at DIV21 and DIV49 (FDR < 0.05, |log2FC| > 0.58), shows synaptic pathway enrichment is driven by upregulated genes in DIV21 and downregulated genes in DIV49.*

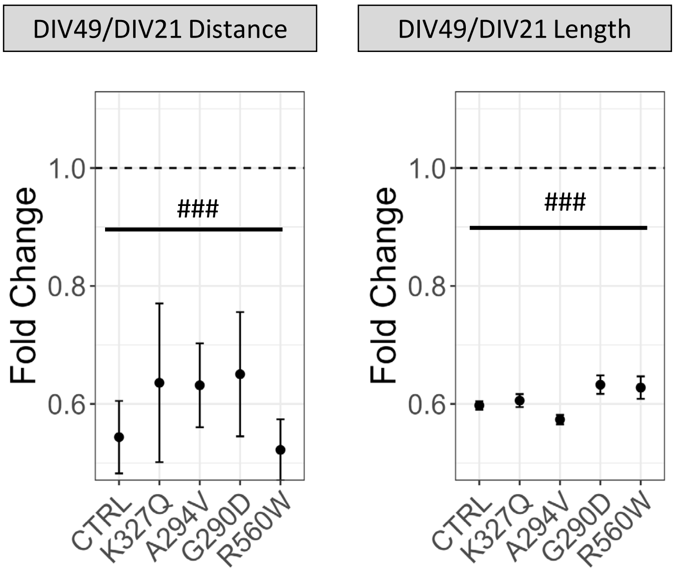


**Fig. S11 Development of the AIS over time.** Graph represents the fold change of AIS distance(left) and length (right) at DIV49 compared to DIV21. Both AIS distance and length shorten over time for all genotypes. Data is presented as median +/- SEM.
