## Supplementary material for "KCNQ2 Loss-of-Function variants disrupt neuronal maturation via early hyperexcitability followed by maladaptive network remodeling": Tables 2-7

| | ISO_G290D | G290D |
|---|---|---|
| Resting membrane potential (mV) | -49.1±1,5 (31) | -50.0±1.4 (42) |
| Input resistance (Ri; MΩ) | 757.6±45.5 (16) | 698.5±67.9 (19) |
| Capacitance (pF) | 18.6±1.2 (35) | 20.4±1.1 (44) |
| Rheobase | 20.5±2.3 (21) | 24.2±2.0 (20) |
| AP Threshold (mV) | -34.3±1.4 (18) | -34.7±0.9 (18) |
| AP Amplitude (mV) | 104.9±3.2 (20) | 113.5±3.5 (20) |
| AP Half-Width (ms) | 2.8±0.2 (21) | 2.7±0.3 (20) |
| fAHP | -54.5±1.3 (21) | -54.3±1.2(20) |

Table 2. Overview of current-clamp generated single AP properties. Data is presented as mean+/-SEM. Amount of cells measured are indicated between brackets. AP: action potential, fAHP: fast afterhyperpolarization.

| Parameters from 80% cut off | Parameters from 90% cut off |
|---|---|
| Burst Duration body (BD body) | Burst Duration total (BD total) |
| BD body CV | BD total CV |
| Spikes per burst body (SPB body) | Spikes per burst total (SPB total) |
| SPB body CV | SPB total CV |
| Burst interspike interval (B_ISI) | non burst ISI (nonB_ISI) |
| B_ISI CV | nonB_ISI CV |
| Burst Frequency (BF) | interburst interval (IBI) |
| Burst Peak (BP) | perc Spikes in Bursts (% spikes_B) |
| BP CV | |

Table 3. Overview of network parameters extracted from a 80% and 95% peak-to-baseline cut-offs.

| Phase 1 | | | |
|---|---|---|---|
| **Gene** | **Avg_Expression** | **gene_names** | **FC KCNQ2:geneX** |
| ENSG00000075043 | 14676.36 | KCNQ2 | 1.00 |
| ENSG00000055118 | 3647.93 | KCNH2 | 4.02 |
| ENSG00000184156 | 3526.16 | KCNQ3 | 4.16 |
| ENSG00000129159 | 2843.09 | KCNC1 | 5.16 |
| ENSG00000171303 | 2499.76 | KCNK3 | 5.87 |
| ENSG00000182132 | 2345.96 | KCNIP1 | 6.26 |
| ENSG00000171385 | 2313.79 | KCND3 | 6.34 |
| ENSG00000177301 | 2194.11 | KCNA2 | 6.69 |
| ENSG00000156113 | 2160.30 | KCNMA1 | 6.79 |
| ENSG00000131398 | 2058.57 | KCNC3 | 7.13 |
| ENSG00000069424 | 1971.01 | KCNAB2 | 7.45 |
| ENSG00000120049 | 1867.56 | KCNIP2 | 7.86 |
| ENSG00000158445 | 1763.34 | KCNB1 | 8.32 |
| ENSG00000105642 | 1627.86 | KCNN1 | 9.02 |
| ENSG00000100433 | 1438.55 | KCNK10 | 10.20 |
| ENSG00000151079 | 1425.87 | KCNA6 | 10.29 |
| ENSG00000116396 | 1369.38 | KCNC4 | 10.72 |
| ENSG00000169427 | 1326.94 | KCNK9 | 11.06 |
| ENSG00000162989 | 1031.10 | KCNJ3 | 14.23 |
| ENSG00000184408 | 939.80 | KCND2 | 15.62 |

| Phase 2 | | | |
|---|---|---|---|
| **Gene** | **Avg_Expression** | **gene_names** | **FC KCNQ2:geneX** |
| ENSG00000075043 | 21535.63 | KCNQ2 | 1.00 |
| ENSG00000184156 | 12043.18 | KCNQ3 | 1.79 |
| ENSG00000177301 | 7085.99 | KCNA2 | 3.04 |
| ENSG00000055118 | 6127.94 | KCNH2 | 3.51 |
| ENSG00000171303 | 5985.69 | KCNK3 | 3.60 |
| ENSG00000116396 | 5811.15 | KCNC4 | 3.71 |
| ENSG00000069424 | 5381.11 | KCNAB2 | 4.00 |
| ENSG00000156113 | 5210.37 | KCNMA1 | 4.13 |
| ENSG00000105642 | 4783.53 | KCNN1 | 4.50 |
| ENSG00000129159 | 4339.28 | KCNC1 | 4.96 |
| ENSG00000182132 | 3996.80 | KCNIP1 | 5.39 |
| ENSG00000171385 | 3266.03 | KCND3 | 6.59 |
| ENSG00000158445 | 3148.26 | KCNB1 | 6.84 |
| ENSG00000131398 | 3129.03 | KCNC3 | 6.88 |
| ENSG00000151079 | 2780.57 | KCNA6 | 7.75 |
| ENSG00000169427 | 2556.06 | KCNK9 | 8.43 |
| ENSG00000184261 | 2476.26 | KCNK12 | 8.70 |
| ENSG00000107147 | 2349.44 | KCNT1 | 9.17 |
| ENSG00000184408 | 1874.52 | KCND2 | 11.49 |
| ENSG00000162989 | 1651.03 | KCNJ3 | 13.04 |

Table 4. Top 10 potassium channel genes expressed during phase 1 and phase 2. table shows ensemble gene ID, average gene expression, gene name and the fold change of KCNQ2 compared to the respective gene.

| Phase 1 | | | | |
|---|---|---|---|---|
| **Gene name** | **Synaptic process** | **Effect** | **Biologically relavent in iNeurons?** | **reference** |
| ALK | Neurite outgrowth | Increase | Yes | 1 |
| BAIAP3 | Vesicle Release | Increase | Yes | 2 |
| BCAN | Synaptic Strength/Stability | Increase | Yes | 3,4 |
| BDNF | Neurite outgrowth | Increase | Yes | 5,6 |
| | Calcium-cAMP singaling | Increase | | |
| | NMDA signaling | Increase | | |
| C1QL2 | Vesicle Release | Increase | Yes | 7 |
| CALCRL | Calcium-cAMP singaling | Increase | Yes | 8,9 |
| CBLN2 | NMDA signaling | Increase | Yes | 10,11 |
| | AMPA signaling | Decrease | | |
| CBLN4 | Synaptic Strength/Stability | Increase | Yes | 10,11 |
| CHRNB3 | Calcium-cAMP singaling | Increase | Yes | 12 |
| CNTN5 | Neurite outgrowth | Increase | Unclear effect on synaptic signaling | 13 |
| DOC2B | Vesicle Release | Increase | Yes | 14 |
| FRMPD4 | Synaptic Strength/Stability | Increase | Yes | 15 |
| GRIK4 | Vesicle Release | Increase | Yes | 16 |
| | AMPA signaling | Increase | | |
| GSG1L | AMPA signaling | Decrease | Yes | 17 |
| HAP1 | AMPA signaling | Increase | Yes | 18 |
| IGSF21 | Inhibitory signaling | Increase | Yes | 19 |
| KCTD12 | Calcium-cAMP singaling | Increase | No, inhibitory | 20 |
| | Inhibitory signaling | Decrease | | |
| KCTD16 | Inhibitory signaling | Decrease | No, inhibitory | 21 |
| LGI1 | Neurite outgrowth | Increase | Yes | 22 |
| | Calcium-cAMP singaling | Decrease | | |
| | AMPA signaling | Increase | | |
| | Synaptic Strength/Stability | Increase | | |
| MGLL | Calcium-cAMP singaling | Increase | Yes | 23,24 |
| NOS1AP | Neurite outgrowth | Decrease | Yes | 25 |
| | NMDA signaling | Unclear | | |
| NR3C2 | Vesicle Release | Increase | No, no Mineralocorticoids present | 26,27 |
| OPRK1 | Calcium-cAMP singaling | Decrease | Yes | 28 |
| RAMP1 | Calcium-cAMP singaling | Increase | Yes | 9 |
| SLC24A2 | Caclium Clearance | Increase | Yes | 29 |
| SYNPR | Vesicle Release | Increase | Yes | 30 |
| SYT17 | Neurite outgrowth | Increase | Yes | 31 |
| | AMPA signaling | Unclear | | |

| Phase 2 | | | | |
|---|---|---|---|---|
| **Gene name** | **Synaptic process** | **Effect** | **Biologically relavent in iNeurons?** | **reference** |
| ADCYAP1 | Calcium-cAMP singaling | Decrease | Yes | 32,33 |
| ADRA1A | Calcium-cAMP singaling | Decrease | Unclear, no presence of catecholamines norepinephrine | 34 |
| CALB1 | Caclium Clearance | Decrease | Yes | 35 |
| CRH | Calcium-cAMP singaling | Decrease | Yes | 36 |
| EPHA7 | Neurite outgrowth | Increase | Unclear effect on synaptic signaling | 37,38 |
| FLRT3 | Synaptic Strength/Stability | Decrease | Yes | 39 |
| GRID2 | NMDA signaling | Decrease | Yes | 11,40 |
| | AMPA signaling | Increase | | |
| | Synaptic Strength/Stability | Decrease | | |
| GRM3 | Calcium-cAMP singaling | Increase | Yes | 41 |
| HTR1B | Calcium-cAMP singaling | Increase | Unclear, no presence of serotonine | 42,43 |
| HTR2A | Calcium-cAMP singaling | Decrease | Unclear, no presence of serotonine | 42,43 |
| IQSEC2 | Vesicle Release | Decrease | Yes | 44,45 |
| | AMPA signaling | Increase | | |
| LAMP5 | Vesicle Release | Decrease | Unclear, no presence of inhibitory neurons | 46 |
| Rab17 | Neurite outgrowth | Decrease | Unclear effect on synaptic signaling | 47 |
| TENM1 | Uncertain | Decrease | Unclear function | 48,49 |
| TRPC5 | Vesicle Release | Decrease | Yes | 50 |
| | Calcium-cAMP singaling | Decrease | | |
| IL1RAP | NMDA signaling | Decrease | Yes | 51 |

Table 5. Functional implications on glutamatergic synaptic function of uniquely enriched up and down regulated DEGs at phase 1 and phase 2 identified with SynGo analysis.

| Primers | Target | |
|---|---|---|
| KCNQ2ex6F | KCNQ2 exon 6 | GTAGGGGAAGAGGAGAGAGGGCT |
| KCNQ2ex6R | KCNQ2 exon 6 | GAACAAGGCCTCTGACCCTGAG |
| KCNQ2ex7F | KCNQ2 exon 7 | TGGGGATATGGGTTCCTGGATTA |
| KCNQ2ex7R | KCNQ2 exon 7 | CAGCACCCACACAAGGCAAG |
| KCNQ2ex15F | KCNQ2 exon 15 | AGAGCAGGGCCACATAAAGGC |
| KCNQ2ex15R | KCNQ2 exon 15 | GGAGAGATGGGAGAGACAGCAGA |
| CLYBL-NGN1,2 F | NGN insertion | GTAAACCACTGTGGGGTGGA |
| CLYBL-NGN1,2 R | NGN insertion | AGCAAAAGACCCGACTCAGA |
| CLYBL Insertion site F | CLYBL locus | CAAACATGGCTCAGTTGTGAA |
| CLYBL Insertion site R | CLYBL locus | TTGGTGGTGGTCACAGTCAT |

Table 6. Overview of primers used in this study

| Target | Marker | Host Species | Clonality | Supplier | Cat. Nr | diltution factor |
|---|---|---|---|---|---|---|
| MAP2 | Neuronal cell body and dendrites | Chicken | Polyclonal | Synaptic Systems | 188 006 | 2000 |
| Homer1 | Excitatory post-synaps | Rabbit | Polyclonal | Synaptic Systems | 160 003 | 1000 |
| Bassoon | Pan-pre-synaps | Mouse | Monoclonal | Antibodies Inc. | 75-491 | 1000 |
| Ankyrin-G | Axon initial segment | Mouse | Monoclonal | Antibodies Inc. | 75-146 | 250 |

Table 7. Overview of antibodies used in this study

1. Defaye M, Iftinca MC, Gadotti VM, Basso L, Abdullah NS, Cumenal M, et al. The neuronal tyrosine kinase receptor ligand ALKAL2 mediates persistent pain. J Clin Invest. 2022;132(12).
2. Zhang X, Jiang S, Mitok KA, Li L, Attie AD, Martin TFJ. BAIAP3, a C2 domain-containing Munc13 protein, controls the fate of dense-core vesicles in neuroendocrine cells. J Cell Biol. 2017;216(7):2151-66.
3. Mueller-Buehl C, Reinhard J, Roll L, Bader V, Winklhofer KF, Faissner A. Brevican, Neurocan, Tenascin-C, and Tenascin-R Act as Important Regulators of the Interplay Between Perineuronal Nets, Synaptic Integrity, Inhibitory Interneurons, and Otx2. Front Cell Dev Biol. 2022;10:886527.
4. Sonntag M, Blosa M, Schmidt S, Reimann K, Blum K, Eckrich T, et al. Synaptic coupling of inner ear sensory cells is controlled by brevican-based extracellular matrix baskets resembling perineuronal nets. BMC Biol. 2018;16(1):99.
5. Carvalho AL, Caldeira MV, Santos SD, Duarte CB. Role of the brain-derived neurotrophic factor at glutamatergic synapses. Br J Pharmacol. 2008;153 Suppl 1(Suppl 1):S310-24.
6. Martin JL, Finsterwald C. Cooperation between BDNF and glutamate in the regulation of synaptic transmission and neuronal development. Commun Integr Biol. 2011;4(1):14-6.
7. Koumoundourou A, Rannap M, De Bruyckere E, Nestel S, Reissner C, Egorov AV, et al. Regulation of hippocampal mossy fiber-CA3 synapse function by a Bcl11b/C1ql2/Nrxn3(25b+) pathway. Elife. 2024;12.
8. Anderson LE, Seybold VS. Calcitonin gene-related peptide regulates gene transcription in primary afferent neurons. J Neurochem. 2004;91(6):1417-29.
9. Russell FA, King R, Smillie SJ, Kodji X, Brain SD. Calcitonin gene-related peptide: physiology and pathophysiology. Physiol Rev. 2014;94(4):1099-142.
10. Dai J, Liakath-Ali K, Golf SR, Sudhof TC. Distinct neurexin-cerebellin complexes control AMPA- and NMDA-receptor responses in a circuit-dependent manner. Elife. 2022;11.
11. Sudhof TC. Cerebellin-neurexin complexes instructing synapse properties. Curr Opin Neurobiol. 2023;81:102727.
12. Shen JX, Yakel JL. Nicotinic acetylcholine receptor-mediated calcium signaling in the nervous system. Acta Pharmacol Sin. 2009;30(6):673-80.
13. Mercati O, Danckaert A, Andre-Leroux G, Bellinzoni M, Gouder L, Watanabe K, et al. Contactin 4, -5 and -6 differentially regulate neuritogenesis while they display identical PTPRG binding sites. Biol Open. 2013;2(3):324-34.
14. Bourgeois-Jaarsma Q, Verhage M, Groffen AJ. Doc2b Ca(2+) binding site mutants enhance synaptic release at rest at the expense of sustained synaptic strength. Sci Rep. 2019;9(1):14408.
15. Lee HW, Choi J, Shin H, Kim K, Yang J, Na M, et al. Preso, a novel PSD-95-interacting FERM and PDZ domain protein that regulates dendritic spine morphogenesis. J Neurosci. 2008;28(53):14546-56.
16. Arora V, Pecoraro V, Aller MI, Roman C, Paternain AV, Lerma J. Increased Grik4 Gene Dosage Causes Imbalanced Circuit Output and Human Disease-Related Behaviors. Cell Rep. 2018;23(13):3827-38.
17. Gu X, Mao X, Lussier MP, Hutchison MA, Zhou L, Hamra FK, et al. GSG1L suppresses AMPA receptor-mediated synaptic transmission and uniquely modulates AMPA receptor kinetics in hippocampal neurons. Nat Commun. 2016;7:10873.
18. Mandal M, Wei J, Zhong P, Cheng J, Duffney LJ, Liu W, et al. Impaired alpha-amino-3-hydroxy-5-methyl-4-isoxazolepropionic acid (AMPA) receptor trafficking and function by mutant huntingtin. J Biol Chem. 2011;286(39):33719-28.
19. Tanabe Y, Naito Y, Vasuta C, Lee AK, Soumounou Y, Linhoff MW, et al. IgSF21 promotes differentiation of inhibitory synapses via binding to neurexin2alpha. Nat Commun. 2017;8(1):408.
20. Cathomas F, Stegen M, Sigrist H, Schmid L, Seifritz E, Gassmann M, et al. Altered emotionality and neuronal excitability in mice lacking KCTD12, an auxiliary subunit of GABAB receptors associated with mood disorders. Transl Psychiatry. 2015;5(2):e510.
21. Ren Y, Liu Y, Zheng S, Luo M. KCTD8 and KCTD12 Facilitate Axonal Expression of GABA(B) Receptors in Habenula Cholinergic Neurons. J Neurosci. 2022;42(9):1648-65.
22. Baudin P, Cousyn L, Navarro V. The LGI1 protein: molecular structure, physiological functions and disruption-related seizures. Cell Mol Life Sci. 2021;79(1):16.
23. Dinh TP, Carpenter D, Leslie FM, Freund TF, Katona I, Sensi SL, et al. Brain monoglyceride lipase participating in endocannabinoid inactivation. Proc Natl Acad Sci U S A. 2002;99(16):10819-24.
24. Costa MA, Fonseca BM, Mendes A, Braga J, Teixeira NA, Correia-da-Silva G. The endocannabinoid 2-arachidonoylglycerol dysregulates the synthesis of proteins by the human syncytiotrophoblast. Biochim Biophys Acta. 2016;1861(3):205-12.
25. Candemir E, Kollert L, Weissflog L, Geis M, Muller A, Post AM, et al. Interaction of NOS1AP with the NOS-I PDZ domain: Implications for schizophrenia-related alterations in dendritic morphology. Eur Neuropsychopharmacol. 2016;26(4):741-55.
26. McCann KE, Lustberg DJ, Shaughnessy EK, Carstens KE, Farris S, Alexander GM, et al. Novel role for mineralocorticoid receptors in control of a neuronal phenotype. Mol Psychiatry. 2021;26(1):350-64.
27. Karst H, Berger S, Turiault M, Tronche F, Schutz G, Joels M. Mineralocorticoid receptors are indispensable for nongenomic modulation of hippocampal glutamate transmission by corticosterone. Proc Natl Acad Sci U S A. 2005;102(52):19204-7.
28. Standifer KM, Pasternak GW. G proteins and opioid receptor-mediated signalling. Cell Signal. 1997;9(3-4):237-48.
29. Jeon D, Yang YM, Jeong MJ, Philipson KD, Rhim H, Shin HS. Enhanced learning and memory in mice lacking Na+/Ca2+ exchanger 2. Neuron. 2003;38(6):965-76.
30. Kwon SE, Chapman ER. Synaptophysin regulates the kinetics of synaptic vesicle endocytosis in central neurons. Neuron. 2011;70(5):847-54.
31. Ruhl DA, Bomba-Warczak E, Watson ET, Bradberry MM, Peterson TA, Basu T, et al. Synaptotagmin 17 controls neurite outgrowth and synaptic physiology via distinct cellular pathways. Nat Commun. 2019;10(1):3532.

32. Miyata A, Arimura A, Dahl RR, Minamino N, Uehara A, Jiang L, et al. Isolation of a novel 38 residue-hypothalamic polypeptide which stimulates adenylate cyclase in pituitary cells. Biochem Biophys Res Commun. 1989;164(1):567-74.
33. Johnson GC, May V, Parsons RL, Hammack SE. Parallel signaling pathways of pituitary adenylate cyclase activating polypeptide (PACAP) regulate several intrinsic ion channels. Ann N Y Acad Sci. 2019;1455(1):105-12.
34. Jiao X, Gonzalez-Cabrera PJ, Xiao L, Bradley ME, Abel PW, Jeffries WB. Tonic inhibitory role for cAMP in alpha(1a)-adrenergic receptor coupling to extracellular signal-regulated kinases 1/2. J Pharmacol Exp Ther. 2002;303(1):247-56.
35. Chard PS, Bleakman D, Christakos S, Fullmer CS, Miller RJ. Calcium buffering properties of calbindin D28k and parvalbumin in rat sensory neurones. J Physiol. 1993;472:341-57.
36. Inda C, Dos Santos Claro PA, Bonfiglio JJ, Senin SA, Maccarrone G, Turck CW, et al. Different cAMP sources are critically involved in G protein-coupled receptor CRHR1 signaling. J Cell Biol. 2016;214(2):181-95.
37. Leonard CE, Baydyuk M, Stepler MA, Burton DA, Donoghue MJ. EphA7 isoforms differentially regulate cortical dendrite development. PLoS One. 2020;15(12):e0231561.
38. Clifford MA, Athar W, Leonard CE, Russo A, Sampognaro PJ, Van der Goes MS, et al. EphA7 signaling guides cortical dendritic development and spine maturation. Proc Natl Acad Sci U S A. 2014;111(13):4994-9.
39. Peregrina C, Del Toro D. FLRTing Neurons in Cortical Migration During Cerebral Cortex Development. Front Cell Dev Biol. 2020;8:578506.
40. Kumagai A, Fujita A, Yokoyama T, Nonobe Y, Hasaba Y, Sasaki T, et al. Altered Actions of Memantine and NMDA-Induced Currents in a New Grid2-Deleted Mouse Line. Genes (Basel). 2014;5(4):1095-114.
41. Niswender CM, Conn PJ. Metabotropic glutamate receptors: physiology, pharmacology, and disease. Annu Rev Pharmacol Toxicol. 2010;50:295-322.
42. Nishijo T, Suzuki E, Momiyama T. Serotonin 5-HT(1A) and 5-HT(1B) receptor-mediated inhibition of glutamatergic transmission onto rat basal forebrain cholinergic neurones. J Physiol. 2022;600(13):3149-67.
43. Berumen LC, Rodriguez A, Miledi R, Garcia-Alcocer G. Serotonin receptors in hippocampus. ScientificWorldJournal. 2012;2012:823493.
44. Rogers EJ, Jada R, Schragenheim-Rozales K, Sah M, Cortes M, Florence M, et al. An IQSEC2 Mutation Associated With Intellectual Disability and Autism Results in Decreased Surface AMPA Receptors. Front Mol Neurosci. 2019;12:43.
45. Tagliatti E, Fadda M, Falace A, Benfenati F, Fassio A. Arf6 regulates the cycling and the readily releasable pool of synaptic vesicles at hippocampal synapse. Elife. 2016;5.
46. Tiveron MC, Beurrier C, Ceni C, Andriambao N, Combes A, Koehl M, et al. LAMP5 Fine-Tunes GABAergic Synaptic Transmission in Defined Circuits of the Mouse Brain. PLoS One. 2016;11(6):e0157052.
47. Mori Y, Fukuda M, Henley JM. Small GTPase Rab17 regulates the surface expression of kainate receptors but not alpha-amino-3-hydroxy-5-methyl-4-isoxazolepropionic acid (AMPA) receptors in hippocampal neurons via dendritic trafficking of Syntaxin-4 protein. J Biol Chem. 2014;289(30):20773-87.
48. Cheung A, Schachermayer G, Biehler A, Wallis A, Missaire M, Hindges R. Teneurin paralogues are able to localise synaptic sites driven by the intracellular domain and have the potential to form cis-heterodimers. Front Neurosci. 2022;16:915149.
49. Zhang X, Lin PY, Liakath-Ali K, Sudhof TC. Author Correction: Teneurins assemble into presynaptic nanoclusters that promote synapse formation via postsynaptic non-teneurin ligands. Nat Commun. 2023;14(1):2957.
50. Schwarz Y, Oleinikov K, Schindeldecker B, Wyatt A, Weissgerber P, Flockerzi V, et al. TRPC channels regulate Ca2+-signaling and short-term plasticity of fast glutamatergic synapses. PLoS Biol. 2019;17(9):e3000445.
51. Gardoni F, Boraso M, Zianni E, Corsini E, Galli CL, Cattabeni F, et al. Distribution of interleukin-1 receptor complex at the synaptic membrane driven by interleukin-1beta and NMDA stimulation. J Neuroinflammation. 2011;8(1):14.
