## Supplementary material for "KCNQ2 Loss-of-Function variants disrupt neuronal maturation via early hyperexcitability followed by maladaptive network remodeling": Table1

| Phenotype | Nucleotide variant | Location | zygosity | Sex | Age iPSC generation (y) | Inheritance | Onset Epilepsy | Age of seizure freedom | Development |
|---|---|---|---|---|---|---|---|---|---|
| SeLFNE | c.979A>C;<br><br>p.K327Q | C-terminus | htz | F | 12 | Inherited from affected father and present in affected sibling | 3rd day of life | 2 m.<br>Isolated seizure at 4 y and 5 y | Normal |
| DEE | c.881C>T;<br><br>p.A294V | Pore loop between S5 and S6 | htz | F | 4 | *de novo* | 1st day of life | not | Severe ID, non-verbal, wheelchair-bound |
| DEE | c.869G>A;<br><br>p.G290D | Pore loop between S5 and S6 | htz | F | 13 | *de novo* | 2nd day of life | 12 y and 8 m | Severe ID, challenging behavior (aggressive, irritable, autistic features), non-verbal, walks independently since age 4y 8m |
| DEE | c.1678C>T;<br><br>p.R560W | C-terminus | htz | M | 9 | *de novo* | 2nd day of life | 7 y | Severe ID with abnormal dystonic posturing, bilateral convergent squint, non-verbal, walks with support |
| CTRL | Healthy sib of R560W (WT) | | NA | M | 13 | NA | NA | NA | NA |
| CTRL | Isogenic control of G290D (ISO_G290D) | | NA | F | NA | NA | NA | NA | NA |

Table 1: Information on patients and controls from whom iPSCs were derived. KCNQ2 transcript used: NM_172107.4. ID: intellectual disability, htz: Heterozygous, NA: not applicable, F: female, M: male, m: months, y: years
